# Frequency and Laminar Profile of Feature Specific Visual Activity Revealed by Interleaved EEG-fMRI

**DOI:** 10.1101/2024.07.31.605816

**Authors:** Tommy Clausner, José P. Marques, René Scheeringa, Mathilde Bonnefond

## Abstract

The role of cortical oscillations in brain function has been extensively debated, resulting in a variety of theoretical frameworks. Using interleaved simultaneous EEG-fMRI, we examined the layer-specific relationship between oscillatory activity and visual processing. We could demonstrate that *γ*-band activity positively correlates with feature specific signals in superficial layers, but we were able to report a deep layer contribution as well. In addition, we could demonstrate that *α*-band power not only correlates negatively with the feature unspecific BOLD signal, but related to feature specific BOLD as well. Lower frequency *α* was predominantly related to feature unspecific superficial layer BOLD, while upper frequency *α* was found to be related to feature specific BOLD in superficial and deep layers. We conclude that the role of *α*-band oscillations extends beyond widespread inhibition and might be involved in active stimulus processing on the level of visual features.

## Introduction

The involvement of brain oscillations in cortical computations and their role in laminar communication channels are highly debated in the neuroscience community. Although the significance of *γ* band oscillations for cortical computations remains uncertain (Hermes et al., 2019; Shirhatti et al., 2022; Dowdall et al., 2023; Schneider et al., 2021), they have been associated with bottom up stimulus feature specific processing (Fries, 2009, 2015). In line with anatomical findings, *γ* band oscillations are primarily associated with neuronal activity in granular and supra-granular layers (Bastos et al., 2012). Hence, nearly all current frameworks predict that *γ* band oscillations should be observed specifically during ongoing stimulus feature processing in supra-granular layers (Fries, 2015; Bonnefond et al., 2017; Bastos et al., 2012).

In contrast, *α* oscillations are thought to reflect a general inhibitory process (Klimesch et al., 2007; Pfurtscheller et al., 1996; Jensen and Mazaheri, 2010) (but see e.g. Bonnefond et al. 2017 or Bonnefond et al. 2024). The prevailing idea is that *α* is modulated at a global scale with low spatial specificity, possibly explaining the lack of *α* modulation observed in some animal studies using local references. However, an intracranial EEG study revealed that *α* band could be associated with surround suppression (Harvey et al., 2013). This suggests that *α* band oscillations can exhibit high spatial specificity and possibly be tightly linked to the activity of neuronal populations processing specific stimulus features in primary visual regions, or at least act with receptive field specificity. The discrepancies between results indicating high versus low spatial specificities led to the assumption that *α* band activity reflects a family of low-frequency oscillations that serve functionally distinct roles. As proposed by Klimesch in the 90s (Klimesch, 1997; Klimesch et al., 1998; Klimesch, 1999), different frequency bands, lower and upper *α* band, could be associated with these distinct roles (see also Rodriguez-Larios et al. 2022; Bonnefond et al. 2017). Another debate relates to the laminar profile of *α*. Animal and human studies have reported that *α* oscillations are mainly linked to feedback-directed signalling, implying the involvement of top-down processes (van Kerkoerle et al., 2014), possibly coordinated with the involvement of the pulvinar (Saalmann et al., 2012). Stimulation of higher-order visual regions have been found to cause *α* band activity in primary visual regions (van Kerkoerle et al., 2014). Accordingly, *α* oscillations have been reported to be the strongest in the deep layers which receive feedback connections from higher-order regions (van Kerkoerle et al., 2014; Spaak et al., 2012; Scheeringa et al., 2016). However, the laminar profile of *α* oscillations remains debated with other studies. Those mostly use current source density approaches, reporting a stronger power in superficial layers (Haegens et al., 2015; Halgren et al., 2019), receiving feedback projections, but also feed-forward input from granular layer generators. Therefore, additional research is needed to better understand the laminar profile of *α* oscillations, possible different sub-bands and their functional and spatial specificities.

Addressing these questions requires a thorough investigation of feature-specific cortical activity on the laminar level, which has traditionally posed a challenge in healthy human participants. Hence, most evidence has been obtained from animal models. Past studies have primarily focused on methodological obstacles (Scheeringa et al., 2016, 2023; Bonaiuto et al., 2021) or have relied on laminar level functional magnetic resonance imaging (fMRI) alone (Kok et al., 2016; Lawrence et al., 2019a; Self et al., 2019; Stephan et al., 2019; Yang et al., 2021). However, fMRI lacks the temporal resolution required and serves only as a proxy for neuronal activity. In contrast, electroencephalography (EEG) and magnetoencephalography (MEG) directly record neuronal activity, but have relatively low spatial resolution because the signals obtained stem from large populations of neurons. Past studies combining high-resolution (laminar-level) fMRI with EEG recordings have successfully addressed these limitations (Scheeringa et al., 2011, 2016). In our study, we employed this approach to investigate, for the first time, feature-specific BOLD signals in the deep (infra-granular), middle (granular), and superficial (supra-granular) cortical layers in primary visual regions. In addition, we not only examined the relationship between EEG and positive BOLD signal deflections but also related EEG power changes to negative BOLD deflections, addressing the link between *α* and inhibition potentially interfering with unwanted information. Negative BOLD signal deflections have received less attention and mostly focused on the default mode network (Hinz et al., 2019; Mayhew et al., 2013; Raichle et al., 2001), however sparse evidence suggests that negative BOLD deflections contribute to understanding ongoing neuronal processing and specifically the inhibition of unwanted information (Tootell et al., 1998; Shmuel et al., 2006). In our analyses, we focused on feature-specific and feature-unspecific BOLD signal changes, highlighting the general importance of studying BOLD signal decreases for functional tasks.

The primary aim of this study was to investigate the relationship between cortical oscillations and feature-specific and feature-unspecific BOLD signals across multiple cortical layers. We employed a visual oddball task with stimuli consisting of two orthogonal gratings offset by 90^◦^ that served as features of interest (left or right oriented gratings). Our results demonstrate that *γ* band activity not only relates to the feature-specific superficial layer BOLD signal (Bonnefond et al., 2017; van Kerkoerle et al., 2014; Scheeringa et al., 2016; Van Kerkoerle et al., 2017) but also to deep layer BOLD activity. Additionally, we found that general modulatory, feature unspecific processes, potentially associated with attention related mechanisms, and feature-specific laminar activation profiles were linked to distinct *α* frequency bands.

This study provides the first direct evidence in healthy human participants that low-frequency oscillations serve multiple purposes in the visual cortex, associated with distinct cortical layer profiles.

## Results

A dataset consisting of 52 right-handed individuals (34 of whom identified as female) between the ages of 18 and 35 (*µ* = 24.0, *σ* = 4.0) was recorded. We only included participants who did not need eye correction (due to practical reason concerning the scanning procedure) and did not have a history of neurological or psychiatric issues and had not undergone neurosurgery. All participants provided informed consent and were monetarily rewarded for their participation. The study received ethical approval from the local ethics committee.

### Behavioural and intermediate results

On average (SD) participant’s response accuracy was 94% (8%) with a false alarm rate of 2% (3%) to non-oddball stimuli and a miss rate of 5% (7%), indicating that participants performed the task adequately well and complied to the task instructions. The average (SD) reaction time was 759 ms (131 ms). See Figure 1 for a graphical representation of the task.

**Figure 1:**
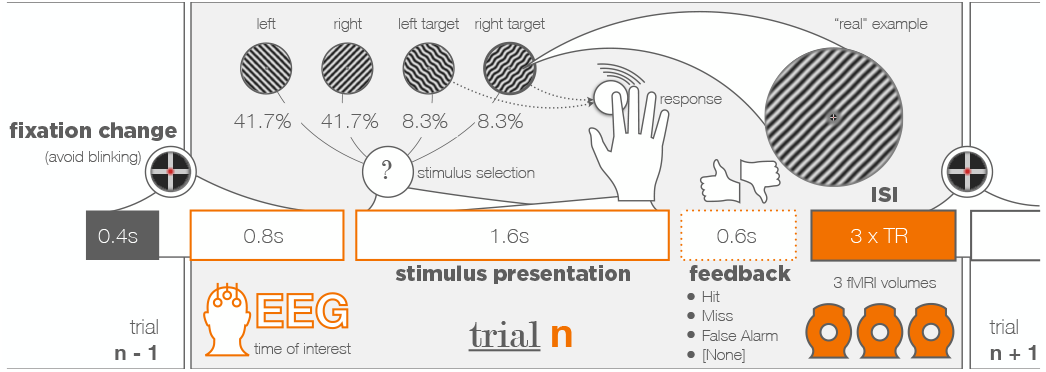
Experimental Procedure of the main experiment. 1.2 s prior to the stimulus onset, the fixation indicator would turn from green to red, indicating the participant to avoid blinking. After a period of 0.4 s (0.8 s before stimulus onset) the last fMRI volume of the previous trial finished recording. For a period of 3 s no fMRI data was recorded in order to avoid gradient artifacts in the EEG data. The stimulus presented is a left or right (*±* 45°from vertical axis) oriented grating. 16.7% of stimuli (2 *×* 8.3% for left and right respectively) were modified to have a slight wavy pattern (see example grating). Participants were asked to respond to those oddball trials with a button press. If no oddball trial was presented, the stimulus would remain on the screen for 1.6 s, followed by a 0.6 s period where only the fixation indicator is shown. In case of a response, corresponding feedback (“correct”, “false alarm” or “miss”) is displayed instead. After that, the fixation indicator turns back to green, indicating to the participant that the period to avoid blinking has ended. Now, three consecutive 3D EPI volumes (TR : 3.3 s) are recorded, before the next trial starts. Overall, 240 trials (four blocks with 60 trials each) have been recorded. Additionally, a high resolution (0.8 mm iso voxel size) T1 weighted full brain image has been recorded before the main experiment to obtain the individual participant’s anatomy. Furthermore, three blocks of pRF mapping (128 trials each) have been performed after the main experiment using the same fMRI sequence as in the main experiment (without gaps).

The EEG signal, used to construct regressors for the final combined EEG-fMRI analyses, was obtained by a time frequency (TF) analysis of each virtual channel, computed for each voxel location of the gray matter in V1. In each hemisphere, one virtual channel for low and one for high frequencies was selected that showed the strongest *α* decrease and *γ* increase respectively. Figure 3 B depicts the average EEG response of the selected, TF transformed virtual channels averaged over all trials and participants. By visual inspection, the main *α* band decrease was determined to last from 0.2 s to 0.8 s after stimulus onset and comprised frequencies from 8 Hz to 14 Hz. The main *γ* band increase was determined between 0.1 s and 0.7 s after stimulus onset for a frequency range between 50 Hz and 70 Hz. A time window between 0.1 s and 0.8 s, was used for the main analyses, since both time windows for *α* and *γ* responses are covered. Furthermore, this time window ensures that only time bins went into the analyses that were related to the actual stimulus processing (*µ*_*RT*_ = 759 ms). To verify that the main TF response of interest originated at the occipital pole, a DICS beamformer full brain analysis had been conducted at the centre frequencies of the selected bands. Figure 3 A depicts the top 5% of the average decrease at 11 Hz or increase at 60 Hz, which localised to occipital regions.

As expected, the strongest BOLD signal increase was found at the occipital pole, due to the central presentation of the stimulus (Engel et al., 1997; Dumoulin and Wandell, 2008). In addition, we observed a clear-cut t-value distribution pattern around the calcarine sulcus, with more negative t-values located dorsal and more positive t-values ventral to the fissure. The first level contrast between both stimulus orientations did not reveal any clear pattern, which served as a sanity check. See Figure 3 C for a visualisation.

### Combined EEG-fMRI analyses

The relationship between the EEG and fMRI data has been investigated on a trial-by-trial basis by means of a general linear model (GLM) as was previously done (Scheeringa et al., 2011, 2016). A separate GLM model was computed for each TF bin by convolving the z-transformed EEG response of either of the four time-frequency transformed virtual channels (2 frequency bands *×* 2 hemispheres) with the standard haemodynamic response function as built into SPM12. Task and nuisance fMRI regressors served as control parameters and were fixed for each model. *β* coefficients for every voxel that were taken into consideration were multiplied by the layer weights for those respective voxels. Before the GLM was computed, the fMRI data was z-transformed across time, separately for each block and voxel. Laminar segmentation was performed by defining five curvature-corrected layers (CSF, superficial, middle, deep, and white matter) between the grey and white matter boundaries, where each voxel received layer weights reflecting the fraction of its volume shared with each respective layer shell (Shafee et al., 2015; Van Mourik et al., 2019a). GLMs were computed for the most activated (or deactivated) 5%, 10% and 25% voxels (only results for the top 10% are presented, see Figures S1–S9 in Supplementary Figures for the full set of results). The data were averaged for a time window between 0.1 s and 0.8 s after stimulus onset. After the GLM, the final cortical depth by frequency matrices of each participant were statistically evaluated by means of a cluster permutation test (Maris and Oostenveld, 2007). Each significant cluster was then averaged along the frequency dimension at the widest point to enable an auto-regressive rank order similarity (aros) test (Clausner and Gentili, 2022), testing the laminar activation profile. The aros test transforms the layer averages into a rank order and tests - using a permutation procedure - if the rank order of the layer averages explains the data better than a random rank order (shuffled layer labels) would. See Figure 2 for a visualisation of the full analysis pipeline.

**Figure 2:**
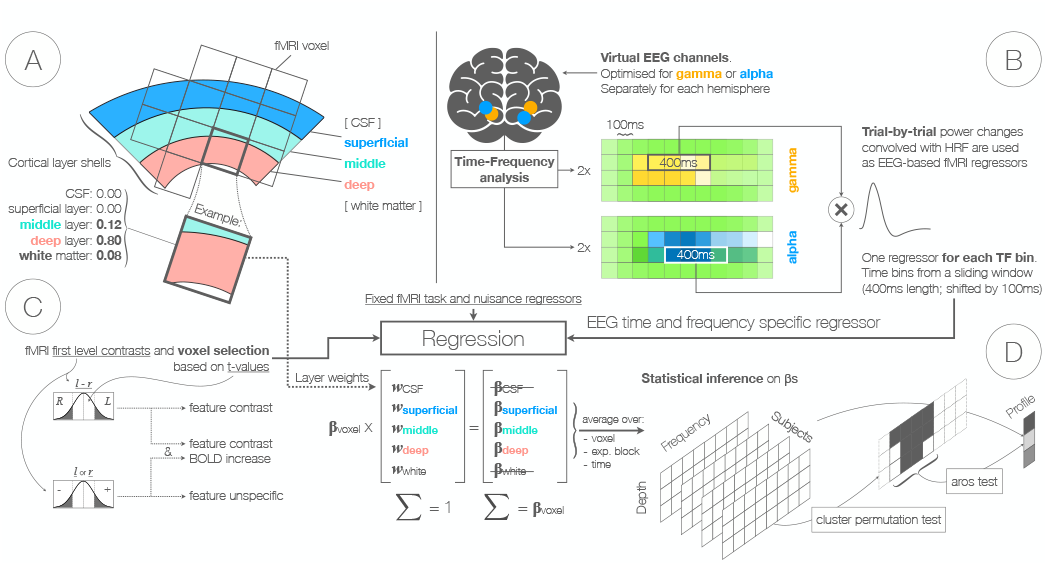
**Data preparation and combined EEG-fMRI analysis** for three cortical layers. **A) Layer specificity**. Cortical layers are constructed as shell-like meshes, taking the gray and white matter boundaries as reference. Between those two reference shells two additional shells divide the space between gray and white matter into three layers. The area outwards relative to the gray matter boundary is assigned to the CSF layer, whereas the area inwards from the white matter boundary is assigned to the white matter layer. Each voxel contributes a fraction of its signal to those layers, depending of the proportional volume of a voxel within each shell like mesh of each layer (see example). Respective fractions are later used as weights to split the *β* coefficients resulting from GLM into different layer contributions. **B) EEG based regressors**. After transforming the EEG data into virtual channel data for each grid point within the gray matter (using LCMV beamforming), virtual channels are time-frequency transformed. For each hemisphere and frequency band separately (2 Hz to 32 Hz for *α* and 20 Hz to 120 Hz for *γ*) the virtual channel with the highest *γ* power increase or *α* power decrease after stimulus onset is selected respectively (∑ = 4 virtual channels). Regressors are built for each time-frequency bin separately. Time bins are averaged to boost SNR. Power values over trials are convolved with the HRF as built into SPM12 resulting in one parameter modulation regressor for each TF bin. **C) Voxel selection**. Voxels are selected based on t-maps resulting from first level contrasts. Thereby, the contrast between both stimulus orientations, and the response to any or both stimulus orientations alone are considered. **D) Statistical inference**. EEG based regressors are entered into a general linear model (GLM) as predictors for the BOLD signal in each selected voxel. The resulting *β* coefficients for each voxel are multiplied with the respective layer weights (excluding white matter and CSF layers) in each voxel and averaged for the respective time window of interest (0.1 s to 0.8 s after stimulus onset) to obtain the final depth by frequency resolved data. This data was tested against the hypothesis that there was no significant relationship between the EEG and fMRI data (*β* coefficients do not differ from zero) for each respective condition, using a cluster permutation test (Maris and Oostenveld, 2007). Resulting clusters were averaged in each layer over frequencies for the widest possible window selected across layers. The layer profiles of the averaged clusters were tested against the hypothesis that the layer profile is as likely as any other layer profile - under the assumption of interchangeability of the data - using an auto-regressive rank order permutation (aros) test (Clausner and Gentili, 2022).

**Figure 3:**
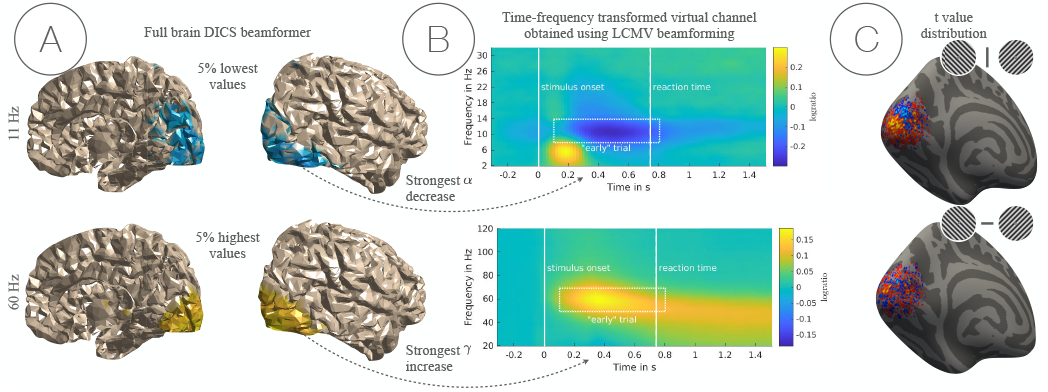
Intermediate results for the EEG and fMRI data. **A) Full brain DICS beamformer results**. Participant average of log-ratios between stimulus and baseline for 11 Hz (*α*) and 60 Hz (*γ*) for both sets of separately filtered EEG data. This serves illustrative purposes only, since the virtual channels of interest have been selected from time-frequency transformed virtual channels obtained using LCMV beamforming. Here, the 5% vertices with the strongest decrease (top) or increase (bottom) are shown. **B) Time-frequency representation of virtual EEG channels**. Participant average of log-ratios between stimulus and baseline of time-frequency transformed virtual channels, obtained using LCMV beamforming (2 *×* 2 virtual channels for low and high frequencies and both hemispheres separately). Only the right hemispheric channels are shown. The white empty square indicates the data points that were included in the combined EEG-fMRI analyses. Average reaction time and stimulus onset are indicated by a continuous or dashed white line, respectively. **C) Average t-value distribution**. Surface projection of average t-map of the first level contrast for the general fMRI activation (top) and the contrast between left and right stimulus orientation (bottom) for illustrative purposes.

During each trial either a *left* or a *right* oriented grating was presented, from which two types of analyses have been derived: *feature unspecific* BOLD activation (i.e. the response to any stimulus orientation), and *feature specific* BOLD activation (i.e. the response to a specific stimulus orientation or the contrast between them). Thereby, fMRI data and EEG based regressors could either be combined congruently (*Co*) by combining the BOLD signal of orientation selective voxels with EEG based regressors built from the same orientation trials, or incongruently (*Inco*), by combining the orientation specific BOLD signal with EEG based regressors built from the other orientation trials. Finally, those two congruency conditions have been contrasted (*Co* − *Inco*).

### Feature unspecific BOLD activation

Feature unspecific BOLD activity was defined as the response to both stimulus orientations combined. In addition to positive feature unspecific BOLD deflection, negative signal changes have been investigated as well. A negative relationship between *α* power changes and BOLD signal change has been found by means of a cluster permutation test (Maris and Oostenveld, 2007). While no difference across layers has been observed for the positive BOLD signal, the strongest negative relationship between *α* and the negative BOLD signal was found in superficial layers. Neither for positive nor for negative voxel sub-selections, a significant relationship between the *γ* band EEG power regressors and the BOLD signal has been observed. See Figure 4 for a visualisation of the results, including p-values corrected using the Benjamini-Hochberg procedure (Benjamini and Hochberg, 1995) to adjust false discovery rates (FDR). We furthermore found a significant interaction (*p*_*FDR*_ *<* 0.05) between positive or negative BOLD signal change and lower or upper *α* sub-bands (8 – 10 or 11 – 13 Hz respectively) by means of a linear mixed effects model. An analysis of simple effects revealed that upper *α* frequencies are stronger negatively related to the positive BOLD signal as compared to lower *α* on a trend level (*p*_*FDR*_ = 0.097).

**Figure 4:**
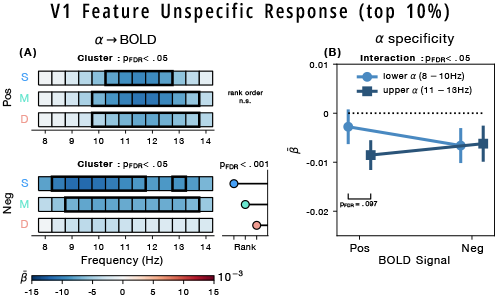
V1 feature-unspecific relationship between laminar BOLD signal and EEG. Average 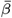 coefficients from a GLM predicting laminar BOLD signals in V1 from trial-by-trial EEG spectral power, shown separately for voxels with a strictly positive (Pos) or strictly negative (Neg) response to both stimulus orientations. 10% most extreme t-values were selected. 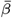 coefficients were weighted by laminar contribution weights, yielding estimates for superficial (**S**), middle (**M**), and deep (**D**) layers. **(A)**: *α*-band (8 – 14 Hz) layer *×* frequency plots. Black rectangles indicate significant clusters (Maris and Oostenveld, 2007); where significant, lollipops to the right show the layer rank order of the effect (Clausner and Gentili, 2022). **(B)**: average 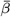 (*±* SEM) for lower (8 – 10 Hz) and upper (11 – 13 Hz) *α* sub-bands per voxel selection, tested with a linear mixed-effects model. All p-values are FDR-corrected (Benjamini and Hochberg, 1995).

### Feature contrast

Contrasting trials where the respective stimulus orientation was *left* with those where the stimulus orientation was *right*, yields the feature specific contrast. Voxels were separated into those with a stronger response to left oriented stimuli (*L*) or those with a stronger response to right oriented stimuli (*R*). Voxel preferences have been combined with EEG power regressors either congruently (*Co*; stimulus orientation of EEG regressor matches voxel preference) or incongruently (*Inco*; stimulus orientation of EEG regressor does not match voxel preference). We found a significant negative relationship between *α* band power and the BOLD signal for *Co* (predominantly in superficial layers) and *Co* − *Inco* (predominantly in superficial and deep layers). A result significant at the trend level (*p*_*FDR*_ = 0.053) for *Inco* was observed as well (predominantly in superficial layers). A linear mixed-effects model was computed to compare upper frequency *α* (11 – 13 Hz) and lower frequency *α* (8 – 10 Hz) with respect to congruency. After correcting for multiple comparisons, we found a significant interaction (*p*_*FDR*_ *<* 0.001). This interaction is mainly driven by the upper *α* sub-band, as indicated by the simple effects analysis. We found a significantly stronger negative relationship of upper *α* and the BOLD signal for congruent selections (*p*_*FDR*_ *<* 0.01) and the reverse for the incongruent condition (*p*_*FDR*_ *<* 0.01), as well as a significantly stronger negative relationship within the upper *α* sub-band for congruent over incongruent voxel selections (*p*_*FDR*_ *<* 0.01). This indicates a more feature related contribution of high frequency *α* and a more general contribution of low frequency *α* to the overall effect. An exploratory analysis of the relationship between individual *α* frequency (IAF) and task performances underlines this finding (see Figure S12 in Supplementary Figures). Thereby the IAF was obtained from the average *α* power spectrum of each participant. The frequency with the strongest decrease between 0.1 and 0.8 s after stimulus onset served as the IAF. We correlated IAF with average response times to correct oddball trials and d’ as a measure for accuracy and found a significant positive correlation between IAF and d’ (*p <* 0.05).

No significant relationship between *γ* band power and the BOLD signal has been observed after correction for multiple comparisons. However, a trend level result has been found for the 25% threshold, with the strongest positive relationship in deep and superficial layers (see Figure S4 i in Supplementary Figures). For a visualisation of the results, including p-values corrected using the Benjamini-Hochberg procedure (Benjamini and Hochberg, 1995) to adjust false discovery rates (FDR), see Figure 5 A-C.

**Figure 5:**
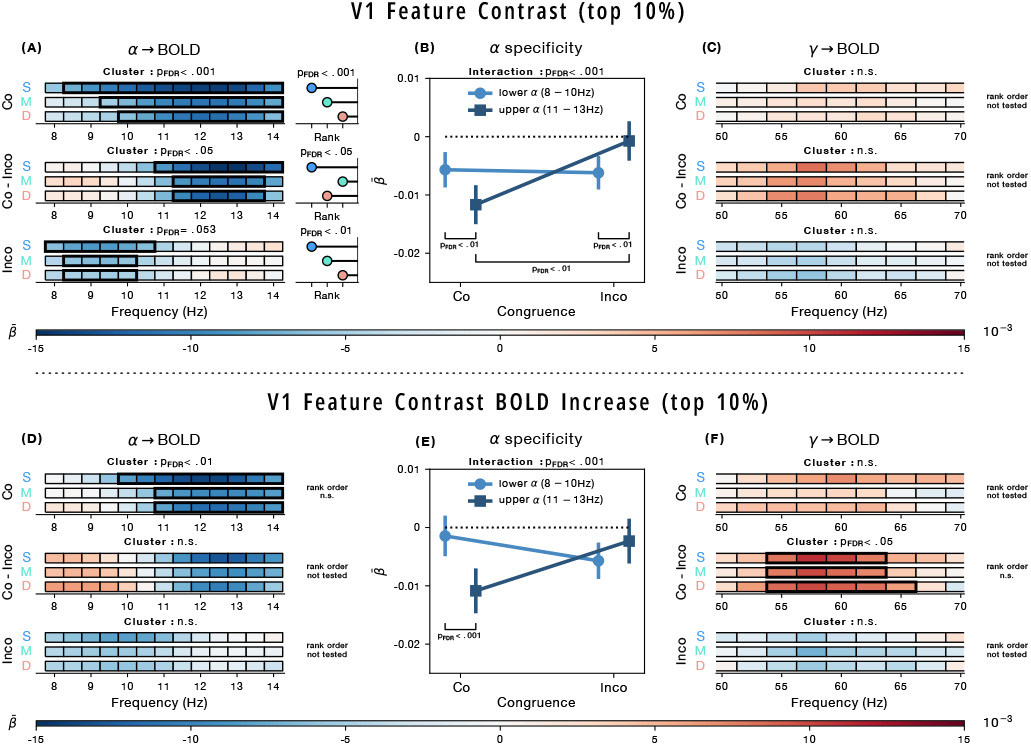
V1 relationship between laminar BOLD signal and EEG for the feature-specific contrast and feature specific BOLD increase. Average 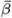 coefficients from a GLM predicting laminar BOLD signals in V1 from trial-by-trial EEG spectral power. Voxels where selected based on the first level fMRI contrast of *left* – *right* oriented stimuli (feature contrast; **A–C**) or with an additional constraint, selecting only voxels with a positive response to any stimulus orientation alone (feature contrast BOLD increase; **D–F**). 10% most extreme t-values were selected. EEG regressors were computed from trials of the Congruent orientation (matching the preferred voxel orientation) or the Incongruent orientation (opposite). Furthermore, the difference between congruent and incongruent models has been computed (Congruent - Incongruent). 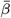 coefficients were weighted by laminar contribution weights, yielding estimates for superficial (**S**), middle (**M**), and deep (**D**) layers. **A, D**: *α*-band (8 – 14 Hz) layer *×* frequency plots. **B, E**: average 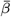 (*±* SEM) for lower (8 – 10 Hz) and upper (11 – 13 Hz) *α* sub-bands per voxel selection, tested with a linear mixed-effects model. **C, F**: *γ*-band (50 – 70 Hz) layer *×* frequency plots. Black rectangles indicate significant clusters (Maris and Oostenveld, 2007); where significant, lollipops to the right show the layer rank order of the effect (Clausner and Gentili, 2022). All p-values are FDR-corrected (Benjamini and Hochberg, 1995). See Figures S4 and S5 in Supplementary Figures for results for the 5%and 25% thresholds.

### Feature specific BOLD increase

The Feature specific BOLD increase was defined similar to the feature specific contrast (see above), but limited to the sub-selection of voxels that respond strictly positive to either orientation. We observed a negative relationship between the *α* frequency band and the feature specific BOLD signal increase for *Co* (*p*_*FDR*_ *<* 0.01). No significant effect was observed for *Inco* or the contrast *Co* – *Inco* in the *α* band. The laminar profile for *Co* has been observed to be flat except for the most liberal threshold (25% most activated voxel) where the strongest relationship between *α* and the BOLD signal was located in superficial layers followed by deep layers (see Figure S4 c in Supplementary Figures). The mixed-effects model for lower frequency *α* (8 – 10 Hz) and upper frequency *α* (11 – 13 Hz) resulted in a significant interaction (*p*_*FDR*_ *<* 0.001) of *α* frequency and congruence condition. This interaction is mainly driven by the upper *α* sub-band, as indicated by the simple effects analysis, which revealed a significantly stronger negative relationship of upper *α* and the BOLD signal for congruent over incongruent selections (*p*_*FDR*_ *<* 0.001). Within the *γ* frequency range, we observed a positive relationship between the *γ* band EEG signal and the feature specific BOLD signal increases for the contrast between congruent and incongruent conditions. Only for the most liberal threshold (25%) a significant layer profile was obtained, with the strongest relationship between *γ* and the BOLD signal in superficial and deep cortical layers (see Figure S5 i in Supplementary Figures). For a visualisation of the results, including p-values corrected using the Benjamini-Hochberg procedure (Benjamini and Hochberg, 1995) to adjust false discovery rates (FDR), see Figure 5 D-F.

## Discussion

In the present study, we combined laminar-level fMRI with simultaneously recorded EEG in healthy human participants to investigate how stimulus-induced changes in *α* and *γ* band power relate to the BOLD signal across cortical layers. We found that *γ* band activity was positively correlated with feature-specific increases of the BOLD signal in superficial and deep layers (see Figure 5). Additionally, we observed a negative relationship between *α* band power and feature-specific BOLD signal activity in superficial and deep layers, primarily for frequencies between 11 and 13 Hz (see Figure 5). Furthermore, we found *α* band oscillations to be negatively related to BOLD signal increases and decreases for *any* stimulus (global modulation; see Figure 4)

The strongest BOLD increase (independent of the stimulus orientation) was found at the occipital pole, which was expected given the central stimulus presentation (Engel et al., 1997; Dumoulin and Wandell, 2008). In addition, we observed reduced BOLD activity dorsal to the calcarine sulcus (Figure 3 C top). Even though attention has not been manipulated as part of this experiment, the overall distribution of positive and negative BOLD reported here, closely resembles previous findings on the retinotopy of the attentional spotlight, where receptive fields at attended locations would exhibit higher BOLD activity and unattended parts lower BOLD activity (Tootell et al., 1998). One interpretation would be that the task could be solved by spatially attending a part of the stimulus only. In addition, negative deflections do probably not reflect a case of “blood stealing” (Smith et al., 2004), since it has been found that a decrease in attention led to a decrease in the BOLD signal (Tomasi et al., 2006), which has been related to decreased neuronal activity (Shmuel et al., 2006).

The combined EEG-fMRI analyses revealed a positive correlation between *γ* and feature-specific voxels. For the most liberal selection threshold (25%, see Figure S5 i in Supplementary Figures) this correlation was observed to be strongest in both superficial and deep layers. The relationship between *γ* and superficial layer BOLD is similar to what has been reported before and was predicted by the literature (Scheeringa et al., 2016; Bonnefond et al., 2017). Previous work in monkeys also attributed *γ* band activity to superficial layers (van Kerkoerle et al., 2014). Accordingly, oscillations in the *γ* band have been attributed mostly to a feed-forward flow of information via superficial anatomical connections (Fries, 2015; Bastos et al., 2012). However, we also found a relationship between *γ* and deep layer BOLD, which was not reported previously. A notable difference to previous work on laminar level EEG-fMRI is that the present study included a contrast between congruent and incongruent features. The tSNR profile (see Figure S10 in Supplementary Figures) does not follow the same pattern as the here reported *γ* band effect. For this reason, we do consider this finding functionally relevant and not related to vascular signal draining (Koopmans et al., 2010). Furthermore, even though van Kerkoerle et al. (2014) discuss the *γ* effect with respect to superficial layer neuronal activity only, their data shows a significant peak in layer 6 as well, which might even be stronger as compared to superficial layers and potentially even stronger modulated by task relevance (see figures 2F and 2G in the respective publication). In fact, an analysis of the relationship between the EEG signal and the BOLD signal that focused on the feature contrast only (*L* – *R*; independent of the response to either stimulus alone) revealed a trend level result with an even stronger deep layer contribution as compared to superficial layers (see Figure S4 i in Supplementary Figures). This strongly implies an involvement of deep layer *γ* oscillations in feature specific processes (Fries, 2009). Recent publications on the information exchange within and between primary visual cortex areas of macaques also reported deep layer *γ* band activity depending on the stimulus material (Gieselmann and Thiele, 2022; Ferro et al., 2021). Those publications challenge the feed-forward exclusivity of *γ* altogether by revealing intra-area feedback communication in V1 from layer 5 to layer 6 and layer 6 to supra-granular layers. Possibly, the relationship between *γ* and deep layer BOLD we observed is also related to similar processes (Vinck et al., 2025). However, it was also shown that feed-forward connections to higher-order regions do exist in deep cortical layers of macaque monkeys too (Markov et al., 2014). Conclusive evidence about the directionality and source of the *γ* related deep layer effect thus remains uncertain. The correlative nature of the here presented approach cannot provide such insights and are likely better addressed in recordings in animal models (van Kerkoerle et al., 2014). Our data however, clearly supports the investigation of feature processing related deep layer *γ* band activity in general. Since *γ* band oscillations are thought to be a marker of ongoing *active* processing of stimuli (Fries, 2009, 2015; Vinck et al., 2023; Spyropoulos et al., 2024) or directly reflect neuronal spiking activity (Ray and Maunsell, 2015), limiting the analysis strictly to voxels with a positive t-value might have increased the SNR substantially, resulting in a stronger effect for feature specific BOLD signal increases as compared to the feature contrast alone. This might explain why the *γ* result for the feature contrast alone did not reach FDR-corrected significance. Based on previous literature, we expected a *γ* band effect for the congruent condition of the feature specific analysis (Scheeringa et al., 2016), which we did not observe. A possible explanation could be the used stimulus material in our experiment as compared to Scheeringa et al. (2016). Muthukumaraswamy and Singh (2013) found that stationary gratings evoke a weaker *γ* band response as compared to moving annular stimuli that have been used by Scheeringa and colleagues. If *γ* is related to the processing of the actual features themselves (e.g. to a column preferably responding to left oriented gratings), then contrasting congruent and incongruent voxel selections provides the largest possible contrast-to-noise ratio (CNR). In turn annular stimuli as previously used might have activated all possible orientations and thus might have greatly boosted *γ* SNR.

Within the *α* band, we found a general negative relationship between *α* band power and the orientation-independent BOLD signal (with a stronger response in superficial layers for the negative BOLD signal; see Figure 4 A). While the negative relationship between *α* band power changes and positive BOLD signal deflections has been reported previously (Zumer et al., 2014; Scheeringa et al., 2011, 2016), a similar relationship with BOLD signal *decreases* has not been studied as extensively. Whether *α* band power increases locally relative to baseline has not been tested. In general, *α* power decreases have been observed very prominently and thus would likely overshadow local increases (see Figure 3 A and B). Hence, we interpret the negative relationship between *α* power and the negative BOLD signal as a reduction of power decreases that would be linked to the locally reduced BOLD activity (Figure 3 C top). The significant interaction between the sign of the BOLD signal deflection and upper or lower *α* bands (see Figure 4 B) further indicates that multiple *α* related processes contribute differentially to positive or negative BOLD. *Active* cortical patches (positive BOLD) are most likely involved in the processing of visual the visual input itself, but also reflect a more general (possibly attention related) activation. In turn the negative BOLD signal might contain less feature specific activation and is most likely related to attention driven deactivation. This hypothesis receives additional support from the trend level difference in *α* sub-bands for positive BOLD, indicating that lower *α* is less related to the active, possibly feature related processes. The absence of this difference for negative BOLD again indicates a broader, more general process. Future experiments manipulating visual features and attention might reveal a differential upper and lower *α* response to attended visual features and a more general relationship between *α* (and possibly superficial layer cortical activity) for suppressed (unattended) receptive fields. This idea would be in line with the hypothesised inhibitory nature of *α* (Klimesch et al., 2007; Jensen and Mazaheri, 2010; Bonnefond et al., 2017; Zumer et al., 2014).

Furthermore, we observed that the relationship between the feature specific BOLD signal and *α* is predominantly linked to frequencies above 11 Hz (see Figure 5 A). An analysis of upper and lower *α* sub-bands revealed a significant interaction between congruence condition and *α* frequency. This interaction was mainly driven by the upper *α* band (11 to 13 Hz). For congruently selected voxels, the negative relationship was significantly stronger (over lower *α*), while for incongruent selection it was significantly weaker. No such difference has been observed for the lower *α* component, which indicates a more feature-specific involvement of upper *α* and a more general modulatory effect for lower *α* frequencies. Since individual frequency variations have been included as a random slope in the linear mixed-effects model, these effects cannot be explained by a subset of participants driving lower or upper *α* separately. Specifically our findings on upper *α* indicate that *α* is not exclusively linked to global signal modulations, which has been the traditional perspective (Klimesch et al., 2007; Jensen and Mazaheri, 2010). More recent theoretical frameworks have predicted that *α* would instead be related to feature specific processes too (Bonnefond et al., 2017, 2024), which would be in line with our findings. Similarly, in a recent opinion article, Stecher et al. (2025) promote the idea of “content-aware” *α*-oscillations. In agreement with our results, the authors argue that *α*-oscillations are related to *content specific* feedback signals, reflected in increased decoding performance based on *α* power of top-down related processes, even prior to the onset of the stimulus (Hetenyi et al., 2025). Accordingly, we interpret the lower *α* effect as a source of general modulation. Thereby, *α* power would possibly decrease in task-relevant pools of neurons and increase in task-irrelevant pools of neurons (i.e. attention-related) to inhibit potentially distracting information (Klimesch et al., 2007; Jensen and Mazaheri, 2010). On the contrary, upper frequency *α* would reflect feature-specific processes, only observed in neurons responding specifically to that stimulus orientation. If the feature congruence contrast was limited to strictly positive voxels (*Co* – *Inco*, Figure 5 D), we did not observe a significant relationship between *α* and the BOLD signal. This could be explained by the limited variability if the analysis focused solely on positive BOLD signal changes, where voxels with a very large variability in response (e.g. strongly positive to one but negative to the other orientation) are excluded. Nevertheless, the differential frequency response, with stronger effects for frequencies between 11 and 13 Hz for congruent voxel selections persists.

The literature on differences in frequency within the *α* band depending on visual stimulus features is sparse. According to Klimesch (1997), lower frequency *α* power was assumed to reflect attentional processes, while upper *α* band power was related to semantic memory demands. Here, lower frequency *α* appears to be stronger modulated with respect to the sign of the BOLD signal change (see Figure 4 B), indicating an involvement in more general processes. This would be in line with the findings on lower frequency *α* reported by Klimesch. In turn, our findings on upper frequency *α* oscillations are more often related to feature specific processes (see Figure 5 B), which could be related to the findings on upper frequency *α* by Klimesch. He related upper frequency *α* power to memory performance under high cognitive load, which in turn could be seen as a proxy for how well the stimuli (i.e. their features) were encoded. A second publication by Rodriguez-Larios et al. (2022) employed a single-participant analysis of independent components, which revealed two dissociable *α* rhythms, both of which were differentially modulated by visual distractors. The lower *α* component increased in power, while the upper frequency *α* component decreased in power under the presence of a visual distractor. Behavioural accuracy was positively related with lower frequency *α* power and negatively related with upper frequency *α*. Again, we interpret those findings as indirectly in line with the here presented results, with the lower frequency *α* being related to more general, possibly attentional processes, and the upper frequency *α* to the content of the memory (i.e. the encoding of the visual stimulus). Furthermore, it could be shown that individual *α* frequency (IAF) and task performance are related, such that higher IAF is linked to higher visual task accuracy (Di Gregorio et al., 2022; Trajkovic et al., 2024; Coldea et al., 2022), potentially via increased perceptual acuity through increased IAF (Tarasi and Romei, 2024). Since behavioural performance in the present study was consistently high at 94% on average and participants were instructed to respond quickly to potential oddball stimuli, a higher *α* frequency might reflect a more successful stimulus encoding and hence faster or more accurate behavioural performance. We exploratively correlated the average IAF during non-oddball trials with the average task accuracy (d’) across participants and indeed found IAF and task performance to be positively correlated (See Figure S12 in Supplementary Figures). This would also be in line with recent theoretical work relating *α* frequency and the speed of visual feature sampling (Bonnefond et al., 2024). An increased “sampling rate” via perceptual windows of opportunity that occur in faster succession, would allow for the extraction of more information in a shorter time window, leading to more accurate responses. In addition, *α* amplitude and frequency have been shown to reflect two distinct processes. While *α* frequency has been related to task performance, amplitude could be demonstrated to be related to visual awareness or confidence judgements about individual task performance (Benwell et al., 2017, 2019; Trajkovic et al., 2024) that could even be shown to depend on the exact cortical region (Samaha et al., 2017). Lastly, higher frequency *α* (extending into the *β* range) has been shown to be causally linked to feed-forward processing, via phase-amplitude coupling (PAC) between feedback related *α* and stimulus induced, feed-forward related *γ*. In a simultaneous transcranial magnetic stimulation (TMS) and EEG study, Trajkovic et al. (2025) demonstrated that brief TMS pulses to the prefrontal cortex in order to activate attentional control, caused increased *γ* band activity over occipital sensor sites through high *α* / *β* PAC with *γ*. This suggests that indeed feedback directed power changes in the upper *α* band support bottom up stimulus processing. Taken together, the here observed stronger effect for upper *α* frequencies for feature specific BOLD is in line with those findings. Whether the here reported difference in *α* frequency indeed reflects computationally different processes, or whether the same process operates at multiple frequencies, remains to be investigated.

Not only did we find a dissociation in the frequency domain between the relationship of *α* and the BOLD signal, but furthermore found that the laminar activation patterns provide further evidence for potentially multiple *α*-related processes. The association between *α* and the BOLD signal was strongest in superficial layers for negative BOLD activity *and* feature specific activity (see Figure 4 A and 5 A). However, deep layer-related *α* effects were limited to feature specific processes only (see Figure 5 A *Co* – *Inco*). These findings suggest that superficial layer *α* reflects a broader, more general process, while deep layer *α* operates more narrowly, linked to the processing of the visual features themselves. Previous findings using laminar fMRI (which did not include the investigation of oscillatory activity), indicate that superficial layer activity might be more related to the modulation of attention (Halgren et al., 2019; Scheeringa et al., 2016; Bastos et al., 2020; Lawrence et al., 2019b), whereas mostly feedback related cortical activity in deep layers has been associated with stimulus features (Kok et al., 2016) and (feature) predictions (Bastos et al., 2020). This would further be in line with research in mice, showing that superficial layer activity acts suppressive on deep layers in order to fine tune stimulus feature selectivity (Pluta et al., 2019). A major concern for laminar fMRI is the vascular draining effect (Markuerkiaga et al., 2016), which typically leads to increased signal amplitudes closer to the surface. Here, we did not investigate the signal change per se, but rather the relationship with EEG power changes. To ensure that the results do not stem from differences in tSNR across layers, we conducted a tSNR analysis (see Figure S10 in Supplementary Figures). We found that the highest tSNR was obtained from deep layers, as compared to middle and superficial layers. To verify, we computed the tSNR using a second layering algorithm (LayNii, see Huber et al. 2021), which yielded lower absolute values, but a comparable layer profile. The obtained tSNR is not reflected in any of our result profiles (see Figures 4 and 5), which strengthens the validity of the here presented results. We speculate that tSNR in deep layers is higher, because both functionally relevant components of the BOLD signal and physiological noise components drain towards superficial layers.

One remaining question would be, how anticipatory and actual stimulus processing compare. It has been suggested that pre-stimulus *α* band activity in the upper range is predictive for task performance (Di Gregorio et al., 2022; Trajkovic et al., 2024), but evidence about the laminar profile of anticipatory processes is still lacking. Due to the limited time window prior to the stimulus onset and gradient artifacts of the MRI machine that lasted until 300 ms before the onset of the stimulus, pre-stimulus *α* could not be investigated in depth here. The visible decrease of *α* prior to the onset of the stimulus (see Figure 3 B) however, indicates that pre-stimulus *α* might play a role for the present task.

A second remaining question is the relationship between the observed *α* and *γ* band effects. For the congruence contrast of feature specific voxels the strongest effects have been observed in superficial layers, followed by deep layers, which again supports the hypothesis that this deep layer effect is at least partly related to feature related processing. If and how lower and upper-frequency *α* band oscillations across different cortical layers interact with each other and how each or both interact with *γ* band oscillations (Bonnefond et al., 2017), needs to be investigated in detail in future experiments. Future work might also vary the exact virtual channel selection for obtaining EEG-based regressors. Here, we focused on the grid points (voxel locations) with the strongest *α* or *γ* response for each frequency band in each hemisphere, derived from the average frequency response to maximise SNR. However, selecting the respective virtual channels based on the response to specific stimulus features or the interaction between high and low frequency bands are possibilities worth exploring in future work.

Furthermore, future experiments might need to consider *β* band activity as well, as those are hypothesised to play a crucial role for visual stimulus processing and might exert a potential top-down influence (Betti et al., 2021; Bastos et al., 2012). Here, we only looked at *β* band oscillations exploratory and found no significant correlation between *β* and the BOLD signal. This could be due to the selection process for the respective EEG virtual channels that were optimised specifically for *α*, but furthermore could be explained by the burst-like nature of *β* oscillations (Bonaiuto et al., 2020; Betti et al., 2021) which makes them harder to capture with the respective analysis strategies employed here.

## Methods and Materials

### Data Acquisition

Functional and anatomical magnetic resonance imaging (fMRI) data were recorded using a Siemens MAGNETOM Prismafit 3T MRI scanner equipped with a 64-channel whole head and neck coil. Before entering the scanner, each participant received detailed instructions and was given the opportunity to practice the main experiment in a short block. Once prepared, the participant was placed inside the scanner and performed an 8 min practice block. A T1-weighted scan was acquired during this time in the sagittal orientation using a 3D MPRAGE sequence (Brant-Zawadzki et al., 1992) with the following parameters: *TR/TI* = 2.2*/*1.1*s*, 11^◦^ flip angle, FOV 256 *×* 256 *×* 180 mm and an 0.8 mm isotropic resolution. Parallel imaging(*iPAT* = 2) was used to accelerate the acquisition, resulting in an acquisition time of 6 min and 31 s. For the functional data, we utilised a 3D gradient-EPI (Poser et al., 2010) with CAIPI acceleration capabilities (Narsude et al., 2016) as implemented by Stirnberg et al. (2017). A partial brain acquisition using a coronal slab was encoded with a FOV 208.8 *×* 208.8 *×* 39.6 mm covering most occipital and parietal lobes, including primary visual regions. The flip angle was set to 20^◦^, resulting in a near isotropic voxel size of 0.9052 *×* 0.9052 *×* 0.9 mm (volume TR: 3.3 s; TE: 34 ms). The sequence was modified to allow an arbitrary time delay between every 3 consecutive volumes. Here, the delay was set to 3 s to ensure unperturbed EEG data acquisition during this delay. The BOLD HRF was subsequently sampled afterwards.

This protocol was used for both, the main experiment and the retinotopic field mapping. For the latter, the sequence gap was omitted because no EEG data was recorded. Each experimental block started with six dummy volumes to allow for the magnetization to reach a steady state, but only the last three of those dummy volumes were actually recorded (and later removed for the data analysis).

In addition, we simultaneously recorded EEG data using a 64 channel MR compatible EEG system (Brain Products Inc, GmbH, Munich, Germany, 2018) at a sampling rate of 5*k* Hz. Impedances were kept below 20*k* Ω during participant preparation. Electrode positions were recorded using a photogrammetry-based approach, as described in Clausner et al. (2017). A 3D model, computed from approximately 50 photographs of each participant wearing an EEG cap, was aligned via facial features to a 3D representation of the anatomical MRI. Electrode positions were determined from the photogrammetry based 3D model, transformed into MRI space and projected along the vertex normals to the MRI scalp surface.

Furthermore, eye-tracking data was simultaneously recorded using an EyeLink 1000+ (SR Research, 2018), but later omitted from the analysis protocol due to insufficient data quality and cumbersome handling in the scanner.

The full experimental protocol comprised a short block of practice trials for the main experiment (8 min), during which the high-resolution anatomical T1 scan was recorded, followed by four consecutive blocks of EEG-fMRI recordings for the main experiment. Each block lasted for 14 min, with a total duration of 4 *×* 14 = 56 min. Three blocks of population receptive field (pRF) mapping were recorded hereafter, each block lasting for 7 min, utilizing the same fMRI recording sequence but without the 3 s pause for clean EEG data recording (3 *×* 7 = 21 min). Additionally, 20 resting-state volumes of that sequence with an inverted flip angle lasting 1 min were acquired for distortion field estimation. However, this was not included in the final analysis protocol. Instead a non-linear recursive boundary estimation (Van Mourik et al., 2019b) was used that simultaneously provides the cortical layer estimation (explained in detail below). The total duration of the experiment was approximately 150 min, including ≈ 40 min preparation time, a 5 – 10 min break between the two main experimental parts, and 15 min for participants to wash and dry their hair after the experiment. Subject motion per block was low, with a mean (SD) framewise displacement (Power et al., 2012) of 0.34 mm (0.24 mm) for the main experiment and 0.23 mm (0.22 mm) for the retinotopy (see also Figure S13 in Supplementary Figures).

### Stimulus presentation

Stimuli were projected onto a screen behind the participant’s head using an EIKI LC XL100 projector (https://www.eiki.com/) with a resolution of 1024 *×* 768 px and a maximum brightness of 5,000 ANSI-lumen, and a contrast ratio of 1000 : 1. The effective field of view comprised a 24 *×* 18° visual angle at a distance of 855 mm relative to the participant’s eyes. Throughout the entire experiment, stimuli were presented in an otherwise dark scanner room. During the anatomical scan, participants were able to read the experiment instructions again, performed another practice block and were asked to remain still with their eyes either opened or closed for the rest of the recording.

### Main experiment

Participants performed a demanding visual attention task, using central stimulus presentation. The stimuli could either be left (counter-clockwise) or right (clockwise) oriented gratings (*±* 45° relative to the vertical axis of the screen). A subtle wavy pattern was incorporated as oddball stimuli and participants were instructed to respond to them using their right index finger. The non-oddball to oddball ratio was set to 5 : 1. Stimuli were presented on a gray background with 50% luminance, using the “Presentation” software (Neurobehavioral Systems, Inc., 2018). A fixation indicator was designed based on the findings of Thaler et al. (2013), which consisted of a black, filled circle overlaid with a white cross (also known as the “Greek cross”), containing a central fixation dot (see Figure 1). This design was found to yield higher fixation performance compared to traditional fixation stimuli, such as simple crosses or dots. In our experiment, the central fixation dot at the centre of the fixation indicator was either red or green, indicating to the participant whether they should avoid blinking (red = avoid blinking).

Feature specific (left or right oriented) stimuli were constructed as Tukey-filtered gratings of 8° visual angle in diameter and a spatial frequency of alternating bright and dark lines of 3.125 cycles per 1° = 25 cycles that were presented at the central screen location. The contrast between bright and dark components was set to 70% luminance change. An area of 0.8° visual angle in diameter was cut out centrally to house the fixation mark. Gratings could be presented in either left or right orientation, deviating *±* 45°from the vertical axis. Additionally, oddball trials were constructed similarly, but with a slightly wavy pattern of an amplitude of 0.3571°visual angle and a frequency of 0.6526 cycles per degree visual angle (≈ 4 cycles across the diameter of the stimulus area). Furthermore four different phase offsets 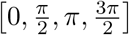 were used in a pseudo-randomised, counter-balanced manner. The outer edge of the stimulus, as well as the edge towards the inner cut-out where the fixation was placed, was filtered using a Tukey filter, to avoid sharp edges.

All settings concerning stimulus appearance were extensively piloted to obtain a satisfactory trade-off between difficulty and accuracy. An example for an oddball stimulus can be found in Figure 1.

### Receptive field mapping

To identify visual cortex regions of interest (ROIs), a population receptive field (pRF) mapping was conducted to obtain structural locations of V1, V2, and V3 for both hemispheres (Arcaro et al., 2009). The experiment consisted of three blocks, with 128 volumes recorded for each block. The stimulus presentation was implemented using the VistaDisp software package (Wandell et al., 2000) in PsychToolbox (Brainard, 1997). A sequence of full-contrast chequered bars was presented, moving in different directions (W → E, SE → NW, N → S, and SW → NE) and their reverse directions in front of an otherwise empty screen of 50% luminance. The bars were 2.25°visual angle wide and up to 18°visual angle long, filling a circular area of 18°visual angle in diameter. The overlap with neighbouring bar locations was 1.125° (half a bar’s width). Sixteen different locations along the directional axis for each moving direction were sampled, resulting in 128 trials per block, with each location being sampled twice. Each location had an alternating full-contrast black and white pattern, presented five times for 0.66 s per cycle (= 0.66 s for 2 alternations). One volume was recorded for one of the sets of five consecutive pattern repetition cycles. For each diagonal moving direction, the pattern disappeared for the last eight locations (40 cycles) of that direction to allow for the BOLD response to fall back to baseline. The procedure is further described in the publication by Alvarez et al. (2015).

### Experimental Procedure

The sequence of events that forms a trial, as well as how the interleaved data acquisition sequence was constructed, is illustrated in Figure 1. After participants were instructed and informed consent was recorded, the EEG cap was fitted, and electrode positions were recorded. Hereafter electrode housings were filled with an electrically conducting gel to bridge the gap between the electrodes and the skull. Afterwards participants were placed inside the scanner. Foam and pillows helped to keep the participant’s head stable and to remain comfortable throughout the experiment. A strap of tape across the forehead provided additional tactile feedback of any head motion and contributed to minimizing head movements. The eye tracking device was set up and calibrated after that. Before the main experiment started - during the high resolution T1 weighted anatomical scan - a practice block was performed for the actual task that followed hereafter. The practice block was a slightly modified version of the main task, such that the inter-stimulus-interval (ISI) was shortened and the ratio of oddball over non-oddball trials was increased to 1 : 3 to facilitate the training effect. The main experiment consisted of four blocks of 60 trials each, ten of which were oddball trials that were excluded from the later analysis. Participants were instructed to respond as fast as possible to the occurrence of such a trial by pressing the response button with their right index finger.

A trial was defined as the following sequence of events: 1.2 s *before* the stimulus onset the central dot of the fixation mark would turn red (indicating the participant to avoid blinking). With a probability of *p* = 0.167 an oddball trial would be shown and otherwise a regular stimulus for 1.6 s. If no oddball was shown and the participant did not respond by a button press, the red fixation stayed for additional 0.6 s before turning back to green, which would end the trial. In all other cases the participant would receive feedback in form of a centrally presented text indicating *hit, miss* or *false alarm*, followed by the green central fixation. Hence, each trial would last 1.2 + 1.6 + 0.6 = 3.4 s of which the last 3 s went into the EEG analysis. During this period, MRI gradients and RF pulses were switched off, such that no MR data could be recorded. This was done to ensure high EEG data quality. After each trial, three partial brain 3D EPI volumes (*TR* = 3.3 s) were recorded, sampling the BOLD response for a single stimulus presentation (Liu and Gao, 2000).

For each of the four experimental blocks, 60 trials were presented, resulting in 240 trials per participant in total. Since trials could be constructed as left or right oriented gratings and oddball or non-oddball trials, four possible trial types could occur. Since the ratio between non-oddball and oddball trials was fixed at 5 : 1, each block consisted of five oddball and 25 non-oddball trials and therefore in total 20 oddball and 100 non-oddball trials for each orientation respectively (∑ = 240). The four different phase-offsets were unevenly distributed within blocks but counterbalanced across the experiment.

After the main experiment, participants could voluntarily rest for some minutes before the population receptive field (pRF) mapping (Dumoulin and Wandell, 2008) was performed. Central fixation during pRF mapping was ensured by a fixation dot that randomly changed colour between red and green at an average rate of 1 change per 3.3 s. This corresponds to 1 TR, since the same fMRI protocol as for the main experiment has been used, only without the gaps that allow noise free EEG recordings during the main experiment. Participants were instructed to indicate a colour change by a button press with their right index finger. All three blocks for pRF mapping were recorded consecutively without a break, except for participant *S*11, where only the first two blocks were recorded due to the participant feeling uncomfortable in the scanner.

### Data processing

Data analyses were performed using the following software packages and toolboxes: analyzePRF (Dumoulin and Wandell, 2008; Kay et al., 2013), ANTs (Avants et al., 2011), FieldTrip (Oostenveld et al., 2011), FreeSurfer (Fischl, 2012), FSL (Smith et al., 2004), janus3D (Clausner et al., 2017), Metashape (Agisoft, St Petersburg, Russia, LLC, 2014), MRICron (Rorden et al., 2012), MRI Volume Masker 3000 TM (Clausner, 2022), MrVista (Wandell et al., 2000), OpenFmriAnalysis (Van Mourik et al., 2019b), SPM12 (Friston, 2007) and Workbench (Washington-University, 2018), including respective dependencies in either Bash, Python or MATLAB.

> The dataset is available at:
>
> https://data.ru.nl/collections/di/dccn/DSC_3018037.01_020
>
> Analysis scripts are available at:
>
> https://github.com/TommyClausner/laminarfMRIv2

No participants have been excluded from any of the analyses. For the main experiment, trials were selected for both non-oddball conditions (= 25 trials per orientation per block = 200 trials per participant). Trials were only excluded if major EEG artifacts such as significant muscle artifacts during the period of interest have been detected, which was the case in ≈ 2% of all trials.

### fMRI Motion Correction and Co-registration

A critical step in laminar fMRI is the correction for motion and co-registration of functional and anatomical data since even sub-millimetre misalignments affect the laminar segmentation substantially. Motion parameter estimation and correction was done using ANTs (Avants et al., 2011). As a first step, the first three volumes were removed from each set of volumes (for the three blocks of pRF mapping and the four blocks of the main experiment). Afterwards a manually drawn brain mask was created for every first volume of the first main experiment block and the first pRF mapping block using MRI Volume Masker 3000 TM (Clausner, 2022). Automatically generated masks were manually “fine tuned”, such that the outer boundary was enclosing the gray matter as closely as possible. Extensive parts of cerebrospinal fluid, fatty components, arteries and other tissue were carefully excluded from the masks. The resulting masks were used to constrain motion parameter estimation and to correct the anatomical segmentation performed by FreeSurfer (Fischl, 2012) within the respective region of interest (the field of view of the functional scans). The actual motion parameter estimation was then performed in two stages. In the first stage all volumes of one recording block were registered to the within-average over time of that block. During the second stage all newly computed within-block averages were registered to the first volume of the first block of the main experiment. Thus all blocks, including pRF mapping, used the first volume of the first block of the main experiment as the final reference. While for the first stage a rigid body transformation was used, an affine transformation was computed for the second stage. The initial linear transformation in that stage was followed by a non-linear transformation using symmetric normalization (SyN, see Avants et al. 2008).

A similar approach was used for functional to anatomical partial volume co-registration, including a rigid body, affine and non-linear transformation using symmetric normalization. Note, that the T1 weighted image was registered to the functional data (and not vice versa) to map the laminar segmentation that is obtained from the anatomical T1 (see below) to functional data space. All estimated motion parameters were combined and applied in a single operation to ensure that functional data was interpolated only once (Esteban et al., 2019).

### Anatomical Segmentation

In a first step, the manually drawn functional masks were registered to native T1 space. This was done in order to ensure proper anatomical segmentation performed using FreeSurfer (Fischl, 2012). This procedure greatly improved the later non-linear boundary registration of pial and white matter boundaries of functional and anatomical data, which are used for the construction of cortical layers and correct for field distortions. In detail, after the initial full brain segmentation, respective functional brain masks obtained as described above were fit to the full brain mask and replaced the respective parietal parts that were covered by the functional data. Hereafter the estimation of pial and white matter surface boundaries was recomputed. Corrected pial surfaces and uncorrected white matter surfaces were used as boundaries for the later laminar segmentation as this procedure makes use of the surface’s boundaries.

### Population Receptive Field Mapping

Population receptive field (pRF) mapping was performed as implemented in the open source toolbox analyzePRF. More detailed information about the algorithmic implementation can be found in the reference literature (Dumoulin and Wandell, 2008; Kay et al., 2013). Binarised versions of each stimulation frame served as spatial regressors for the underlying general linear model (GLM). Each of the presented 64 unique bar locations (including blanks) were thresholded such that the background received a value of 0 and the entire bar irrespective of the chequerboard pattern received a value of 1. In order to save computation time, stimuli were downsampled from screen resolution to a resolution of 192 *×* 192 *px*. A Savitzky–Golay filter with a filter window of 61 TR s (201 seconds) was applied to the data. The data then was converted into percent signal change relative to the median. Whereas the Savitzky–Golay filter was applied for each experimental block separately, percent signal change was computed over all blocks combined. Based on a GLM - including third order polynomials - parameters were estimated for orientation (angle), distance to the centre of the screen (eccentricity) and the explained variance per voxel (*R*^2^). The gray matter mask, obtained from the anatomical segmentation, was applied and only gray matter voxel locations were fed into the pRF analysis. Based on those maps, regions of interest (V1-3) were manually labelled using Freeview. To facilitate the manual drawing process, a functional atlas (Wang et al., 2015) containing all regions of interest was anatomically fitted to the functional data beforehand. Fitted regions from the atlas were overlaid together with the results of the pRF mapping onto the inflated pial surface as obtained from FreeSurfer. Marked labels were then transformed into volumetric data and into functional data space using previously computed co-registration transformation matrices and volumes. It should be noted however, that regions V2 and V3 could not be reliably separated in all participants due to poor data quality. Hence, all analyses will focus on V1 only, but results for V2 and V3 are provided within the Supplementary Figures (see Figures S1–S9).

### Estimation of Cortical Layers

Laminar segmentation was performed using co-registered gray and white matter boundaries as references for upper and lower bounds of the segmentation. In order to resolve cortical depth precisely, the curvature of the anatomical boundaries was taken into account. This is necessary since the relative thickness of cortical layers varies depending on the cortical curvature (Shafee et al., 2015). Each voxel covered by the gray matter mask, received a weight as a function of its volume belonging to each of the shell-like meshes forming the boundaries. If a layer boundary would cut the voxel exactly in half, adjacent layers would receive a weight of 0.5 each. Hence, voxels were not separately treated as belonging to different layers, but rather their signal was seen as a weighted mixture coming from different layers. Thus, a voxel located towards the white matter boundary would contribute more to the signal generated in deeper layers - receiving a higher weight - as compared to a voxel being closer to the surface, which would receive a lower weight at the reference location (Van Mourik et al., 2019a). Layer weights were computed using the open source toolbox OpenFmriAnalysis. As a result, five layer weights per voxel were obtained (CSF, superficial, middle, deep and white matter layer). See also Figure 2 A for a conceptual visualisation and Figure S11 (in Supplementary Figures) for an example.

### EEG data processing

The major goal of the EEG data preprocessing was to optimise noise suppression for each frequency band of interest (*α, γ*) in order to extract EEG signal components of interest as clean as possible. In previously published literature a supervised signal decomposition based on ICA has been used (Scheeringa et al., 2011, 2016). This approach requires the manual selection of target components for each frequency band. By removing all non-target components, noise can be suppressed and the resulting signal will only contain the data of interest. However, beamforming offers the major advantage of being able to perform unsupervised noise suppression (Veen and Buckley, 1988; Adjamian et al., 2009), crucial for extracting the target signal. During the piloting phase, beamformer methods were successfully applied to the EEG data recorded in an (f)MRI environment and yielded high quality source level signals, in line with previous studies using a similar approach (Brookes et al., 2008, 2009).

Low (*α*) and high (*γ*) frequency bands were processed separately to extract the desired response patterns. The data was filtered for the lower frequencies using a pass band between 2 and 32 Hz and for the high frequencies between 20 and 120 Hz respectively. A 50 Hz dft filter to suppress power line noise was applied to the latter as well. EEG electrode locations were obtained from the photogrammetry based 3D model and co-registered using the face shape to the anatomical MRI using janus3D, as described in Clausner et al. (2017). A finite element model (FEM) was computed from the high resolution anatomical T1 based on the FieldTrip-SimBio pipeline (Vorwerk et al., 2018). The leadfield was computed from the EEG electrode positions and the FEM model. Sources were modelled as equivalent current dipoles at locations limited to the respective coordinates of voxels included in the gray matter and ROI masks (e.g. V1). Dipole orientations were derived from the cortical curvature and thickness, since this is crucial for a precise mapping especially in EEG (Baillet et al., 2001). Workbench (Washington-University, 2018) was used to compute the surface normals that connect pial and white matter surfaces. The orientation of the resulting vectors then served as the dipole orientation for each respective location. The described procedure was done separately for the left and right hemisphere in order to obtain separate filter weights, since distinct source activity for both hemispheres was to be expected (Michel et al., 2001). The resulting weight matrices were applied to the band pass filtered data in order to obtain virtual channels at the corresponding equivalent current dipole locations. A spectral analysis was performed on each virtual channel separately. Thereby the exact settings for low and high (time-) frequency decomposition varied slightly. While for low frequencies, a Hanning taper was applied using a data time window length of 400 ms (time steps: 20 ms), zero-padded to achieve a frequency resolution of 0.5 Hz and smoothed with a kernel width of *±* 2 Hz; for high frequencies the data was processed using the same time related settings, applying a multi-taper (DPSS; Slepian 1978) approach (seven tapers) with a resolution of 2.5 Hz. The frequency domain was smoothed using a kernel width of *±* 10 Hz. Afterwards the virtual channel with the highest average amplitude change between 8 – 12 Hz (*α*) or 50 – 70 Hz (*γ*) relative to baseline, was selected (see visualisation in Figure 3 B). As a baseline period, a time bins between −0.3 to −0.1 s for high and a single time bin at −0.3 for low frequencies were chosen. Gradient artifacts caused by ringing of MRI machine’s amplifier after the gradient coils were switched off were observed prior to −0.3 s relative to stimulus onset which was unexpected and not observed during the piloting phase. The initially planned start of the baseline period was −0.5 s, however due to those artifacts −0.3s relative to stimulus onset served as the new lower bound for the baseline period. For this reason, the low frequency baseline period comprised only a single 400 ms time bin centred around −0.3 s, because a pre-stimulus *α* decrease was expected starting around 0.25 s prior to stimulus onset. Pre-stimulus *α* has often been related to visual detection performance (Van Diepen et al., 2019) and might thus reflect different processes than the actual target time interval. In fact a small but visible *α* decrease could be observed 0.25 s prior to stimulus onset (see Figure 3 B). The data was transformed into the log_10_ ratio between the baseline and each sample point in a time window between 0.1 and 1.6 s after stimulus onset for each hemisphere. The time-frequency (TF) transformed virtual channels with the highest average response were chosen to be the “best” channels that were later used to build the regressors for the combined EEG-fMRI analyses. In total four dipole locations (i.e. the TF transform of corresponding virtual channels) were selected for each participant individually: One for each of the two hemispheres and one for each of the two separate frequency bands. Figure 3 A depicts the result of a full-brain DICS beamformer analysis (Gross et al., 2001) for illustrative purposes. Since the LCMV beamformer approach was limited to specific ROIs obtained from pRF mapping, the DICS full brain scan was performed in order to verify that indeed visually induced activity yields the strongest effects at the occipital pole. Note that the main goal was not to accurately localise the source of *α* or *γ* oscillations, but rather to utilise the beamformer’s noise suppressing properties to extract high SNR *α* or *γ* signals programmatically, as opposed to the manual selection of independent components in previous studies (see e.g. Scheeringa et al. 2016). All steps previously mentioned were implemented in MATLAB R2021a (MathWorks, 2021) using the open source toolbox FieldTrip. See Figure 3 B for a depiction of the average TF transformed virtual channel response over participants for one hemisphere and both frequency bands. Since the main hypotheses of the presented study do not directly address the frequency responses (i.e. power changes) themselves, which are well established response patterns, but rather their relation to the BOLD signal, a pre-selection of EEG data with the strongest response, still acts as valid scientific strategy and does not result in a case of “double dipping”.

### Combined EEG-fMRI analysis

The general logic of fitting trial-by-trial based EEG time-frequency regressors individually for each TF bin to the BOLD signal for different cortical regions and across layers, follows largely what is described in Scheeringa et al. (2016). Several steps have been undertaken to prepare the EEG and fMRI data for the later combined analysis.

**Nuisance regressors** contained all trial and response combinations that were not included in the task regressors (e.g. false alarm trials).

Additional regressors contained blink or artifact trials, button presses and reaction times. All the aforementioned regressors were convolved with the haemodynamic response function as built into SPM12. Reaction time regressors were treated as a parametric modulation. The onset of the modulation was set to the average reaction time for each individual block and the actual reaction time as the modulating value. This procedure was chosen in accordance with previous literature (Scheeringa et al., 2016). Furthermore the average white matter signal and the average residual signal (average signal after regressing out gray and white matter signals) were included in the nuisance regressor matrix. All motion parameters (translation along and rotation around x, y, z) and their first derivatives were included as well as a set of high pass filters modelled as five sines and five cosines. Those five sine and cosine waves were constructed, such that they would span one to five full cycles across one experimental block. **Task related fMRI regressors** were built separately for left and right oriented gratings or both combined.

**EEG data regressors** were built on the TF resolved virtual channel data, obtained as described above. Frequency bin span 0.5 Hz for low and 2.5 Hz for high frequency data and time bins were set to 0.4 s that were shifted by 0.1 s intervals (sliding window). One regressor was built for each frequency and time bin separately. This was done by convolving z-transformed trial-by-trial power changes of the EEG data with the haemodynamic response function that comes with SPM12, such that each regressor served as parameter modulator. Thereby the time onset was set to the mid data point of the time domain data bin and shifted by one to three TRs matching the corresponding three volumes that have been recorded after the corresponding trials. In the later general linear model (GLM) each TF based regressor was fit in a separate model. Task and nuisance fMRI regressors however were kept fixed for each model. Similar to fMRI based task-regressors, three sets of task-related regressors were built for left or right oriented trials and both combined. All analyses were performed for a (EEG) time window ranging from 0.1 s to 0.8 s after stimulus onset, for which corresponding time bins were averaged. This was done because the average reaction time to oddball stimuli was 759 ms. Since the main stimulus processing is assumed to take place before the response - but in order to include as many data points as possible - the time bin including 0.8 s after stimulus onset was included as well.

Before the combined EEG-fMRI analysis could be conducted, a normalization step has been applied in order to counteract the signal drop-off resulting from the interleaved sequence. Since in each trial the gap was followed by the collection of 3 consecutive volumes, each voxel of such volumes within one block was divided by the average BOLD signal in that voxel of every 1^*st*^, 2^*nd*^ or 3^*rd*^ volume in that block. Furthermore, a first level fMRI analysis on the motion corrected data was performed in order to identify **voxels of interest**. Thereby multiple selection criteria were applied to the first level t-maps: The *feature unspecific* BOLD response is defined as the response to any stimulus (irrespective of orientation). The *feature specific* BOLD response in turn is defined as the response of a voxel to a specific stimulus over the other (*left* – *right* stimulus orientation). A positive t-value from the contrast thereby indicates a stronger response to left over right oriented gratings, whereas a negative t-value reflects a voxel’s response preference to right oriented gratings. In addition, voxels selected from the feature specific contrast were sub-selected to only include voxels with a strictly positive BOLD response to *any* stimulus (*feature specific BOLD increase*). The threshold used for voxel selections was inconsistent in previous fMRI-EEG publications, for which reason we decided to adopt a transparent approach. Each selection was made such that the top 5%, 10% and 25% of voxels (according to t-value) were included. We corrected for multiple comparisons accordingly. See Figure 2 for a visual representation of the analysis strategy.

Irrespective of voxel selection, a higher **t-value selection threshold** indicates higher specificity for the respective stimulus and a lower threshold increases the signal-to-noise ratio (SNR) by including more data. This changes the number of voxels selected. In previous publications, employing a similar experimental setup, 500 voxels with highest activation (Lawrence et al., 2018, 2019b) or the top 5%, 10% or 25% activated voxels (Scheeringa et al., 2016) were selected. A study by Markuerkiaga et al. (2021), specifically designed to assess the number of voxels required for optimal SNR in the context of laminar level fMRI, finds 250 voxels (for 3T fMRI) to yield the best contrast-to-noise ratio (CNR). However, since all previously mentioned publications set a more or less arbitrary threshold and the last did not take the correlation with EEG into account, three thresholds have been selected to eliminate eventual uncertainties: 5%, 10% and 25% of most activated voxels. We present only the results for the 10% voxel selection threshold, but the full results can be found in Supplementary Figures. In general, results are mostly similar irrespective of the selection threshold.

Importantly, fMRI based voxel selection and EEG based trial-type related regressors could either be combined congruently or incongruently (since feature specific response and the feature contrast can be divided into responding stronger to the left or right oriented stimuli). This means that results could be selected for data of all voxels that respond feature specific to one orientation with EEG based regressors built on trials of exactly this orientation (congruent; short: *Co*) or the respective other orientation (incongruent; short: *Inco*). A congruent pairing (*Co*) would e.g. combine data of each voxel with a stronger BOLD increase to left with EEG based regressors based on trials where the stimulus orientation was left. In the case of *Inco*, the same feature specific voxels would rather be combined in a selection with EEG-based regressors for right oriented stimuli. If voxels for both orientations have been selected (feature unspecific response) an EEG regressor that was built on both trial types was used.

A **general linear model** (GLM) has been computed with predictors for each TF bin separately for all voxels in V1 that later have been sub-selected according to the respective condition. Time courses for each voxel have been z-transformed before the GLM was computed for each voxel and experimental block separately. Afterwards, each of the resulting regression coefficients (*β* values) were multiplied with the voxel specific layer weights that have been obtained as described above. A value of *β >* 0 means that an increase in predictor value (e.g. power of a frequency) correlates with an increase in BOLD (and vice versa). The general strategy rests on the idea that the predicted fMRI activity can be seen as a mixture of signal contributions from each layer in each voxel. For this analysis, contributions from white matter and CSF layers have been excluded and only signal contributions from superficial, middle and deep layers have been taken into account. After averaging results from different hemispheres, experimental blocks, percentiles of interest (e.g. the top 10% voxels of a respective contrast) and time bins for a time interval between 0.1 and 0.8 s after stimulus onset, the final data contains the average regression coefficients for *depth × frequency × participant*. The described procedure of computing a GLM based on z-transformed data using z-transformed predictors yields *β*-coefficients that reflect the average relationship of a single voxel’s BOLD response for a given layer (fraction of the single voxel’s *β*) with EEG power changes of a specified frequency.

Subsequently, separate analyses were done for two frequency of interest (FOI) ranges centred around the *α* and *γ* band responses observed in the EEG. Within these frequency ranges **inferential statistics** based on a cluster level approach were computed. For low frequencies the FOI range was set to frequencies between 8 Hz and 14 Hz (*α* band), whereas for the high frequencies the FOI was defined as frequencies between 50 Hz and 70 Hz (*γ* band), which covers the peak response frequencies found in the average EEG data (see Figure 3 B). Within the respective range a single tailed cluster permutation test (Maris and Oostenveld, 2007) has been conducted separately for each threshold and FOI for *α* and *γ* in V1 since the expected direction of the effect was derived from the literature (Scheeringa et al., 2016; Zumer et al., 2014; Murta et al., 2015). Each significant cluster has been further processed by means of an auto-regressive rank order similarity (aros) test (Clausner and Gentili, 2022). The fundamental idea behind the aros test is whether group averages (i.e. averages of the signal of cortical layer in the present case), can be ranked such that the rank order is explained significantly better by the data than it would if the average data could not be meaningfully sorted (i.e. is shuffled). This is achieved by transforming the group averages into unique rank order values (e.g. superficial *>* deep *>* middle) and computing the average fit of the data to this rank order. In a second step data points are shuffled between the groups and the same procedure is applied (i.e. computing the rank order of the mean and the average fit of the now shuffled data to the new rank order). Repeating this permutation step many times (here 25000 times) yields a permutation distribution, to which the initially computed fit value of the un-shuffled data is compared. Rejecting the null hypothesis would result in the assumption that the rank order of the group averages indeed can be explained by the data significantly better than it would if the data points could not be meaningfully sorted into those groups. Thus, in the present case it could reveal how the correspondence between the EEG and fMRI signals could be sorted across layers. However, statements about the magnitude of the difference between two (or more) layers cannot be made. This approach provides insight about the specific activation profile across layers for specific conditions within a significant *depth × frequency* cluster. Since clusters span layer and frequency bins unevenly, the data that was used for the later aros test was collapsed over frequencies, such that the lowest and highest significant frequency bin served as the boundary over which the frequency domain was averaged. This was done irrespective of the respective cluster size within a specific layer (see Figure 2 D). Testing for layer specificity is not straightforward due to issues related to multiple comparison and non-normal distributed data. To circumvent this, Scheeringa et al. (2016) tested for EEG-fMRI layer specificity by fitting layer profiles for *α* and *γ* to each other using an ordinary regression and tested whether *β* coefficients differed to zero. While this approach is suitable for demonstration purposes, it does not reveal the exact nature of those differences between layers. While Scheeringa et al. proved the concept of layer specific feature extraction, the present paper aims to determine the relational activity across layers depending on the feature specific response as well for which reason the aros test has been developed. For the present case this means that while neglecting the effect size of the difference across layers, the overall profile of the differential activation across layers can be obtained without compromising statistical power due to multiple comparison.

The described procedure of selecting the respective voxels of interest, fitting EEG based regressors jointly with task and nuisance fMRI regressors to the data, weighting the resulting *β* regression coefficients with the corresponding layer contribution weights and applying a cluster permutation test, followed by an aros test on significant clusters, has been applied to each of the contrasts of interest. To account for multiple comparison across the respective selection thresholds, cluster and aros p-values have been adjusted using the false discovery rate (FDR) adjustment procedure proposed proposed by Benjamini and Hochberg (1995).

### Feature unspecific BOLD activation

While previous literature mostly focused on the relationship between BOLD signal increases and different EEG frequency bands, we extended our analysis to include visual cortex BOLD signal *decreases*. In general, the sparse corpus of literature investigating negative BOLD signal deflections in conjunction with the presentation of visual stimuli, mainly focused on attention related effects (Tootell et al., 1998; Tomasi et al., 2006) (especially for foveal presented stimuli and a demanding task; see Crespi et al. 2011). Furthermore, it has been related to decreased neuronal activity in monkey V1 (Shmuel et al., 2006). Previous literature suggests that attention related *α* power increases could be associated with BOLD-signal decreases (Zumer et al., 2014; Shmuel et al., 2006; Tootell et al., 1998) and hence might provide a meaningful insight into feature unspecific signal contributions. Note however that attention was not manipulated as part of this experiment. As described above, for each contrast, multiple thresholds have been applied for voxels with the 5%, 10% and 25% most positive and most negative t-values as a response to any stimulus, of which only the 10% threshold is shown with 5% and 25% in Supplementary Figures.

### Feature specific BOLD response

Previous literature on the relationship between laminar level fMRI and EEG in the visual domain mainly focused on proofs of concept and related visual cortex fMRI activity to EEG without considering different stimulus features (Scheeringa et al., 2011, 2016), which has been accomplished here. In general, the response to the two orthogonal stimulus features has been assessed two fold, using the feature contrast and the feature contrast for voxels with a strictly positive BOLD response. The **feature contrast** is defined by the t-values obtained from the first level analysis contrasting both stimulus orientations (*left* – *right* stimulus orientation). The **feature contrast BOLD increase** only includes voxels from the feature contrast with a positive t-value to any stimulus orientation alone.

Each selection can be separated into voxels that respond preferentially to *left* or *right* stimuli. In addition, the EEG regressors can as well be separated into *left* and *right* by the trials they were constructed from. Using this strategy, the result of the GLM can be separated into congruent or incongruent parts, where feature specific signal changes were assessed for congruent (*Co*) or incongruent (*Inco*) combinations of selected voxels and EEG based regressors. Beside separately looking at congruent and incongruent combinations, the differences between both have been computed as well (*Co* – *Inco*). Thereby, the underlying set of voxels for *Co* and *Inco* is identical and each condition differs only with respect to the EEG regressors used for the GLM. This means *Co* – *Inco* represents differences in estimated response amplitudes between regressors.

Since both stimulus types are exactly orthogonally oriented to each other, the congruence contrast reveals how the relationship of a voxel’s preferred orientation compares to its least preferred orientation, which boosts the feature specific contrast-to-noise ratio. Additionally, the contrast eliminates other external biases (e.g. related to SNR or cognitive processes like attention that have not been manipulated). Each set of results was followed by a single tailed cluster permutation test (Maris and Oostenveld, 2007). The final results structure encompasses the layer *×* frequency resolved data for each voxel selection threshold and combination with EEG power regressors, as well as significant clusters and aros results for *α* and *γ* separately.

### Lower and upper alpha

It has been proposed that *α* band activity is composed of at least two distinct sub-bands with diverging functional roles and neural generators (Sokoliuk et al., 2019; Rodriguez-Larios et al., 2022; Zhao and Liu, 2025). To investigate whether different *α* sub-bands contribute differentially to the negative relationship with the BOLD signal, we additionally conducted multiple analyses for lower (8 – 10 Hz) and upper (11 – 13 Hz) *α* bands. These bands were defined using a data-driven approach. After averaging *α* power over the time window of interest (0.1 – 0.8 s after stimulus onset), we identified the peak frequency (the frequency showing the largest power decrease within the broad *α* range (8 – 14 Hz). We then used the median power across the spectrum as a reference level. The two frequencies at which the power was closest to this median — one below and one above the peak — were taken as the centre frequencies for the lower and upper *α* bands respectively. Finally, a *±* 1 Hz range was applied around each centre frequency, yielding a lower *α* band of 8 – 10 Hz and an upper *α* band of 11 – 13 Hz. *β* coefficients were averaged within those bands and fed into linear mixed-effects models (LMMs) to evaluate the interaction between *α* frequency and experimental conditions. Critically, we employed a unified model structure for both feature-specific (Frequency *×* Congruency) and feature-unspecific (Frequency *×* Activation) analyses. Cortical layers were included as a fixed-effect. Participant-specific variance was modelled using a random-effects structure, including random intercepts for participants and random slopes for frequency and condition (Activation or Congruency). This ensures that observed effects are not driven by individual differences in e.g. *α* frequency. Significant interactions were decomposed into simple effects using Wald tests on the model coefficients, ensuring that post-hoc comparisons were derived from the same statistical global variance as the primary interaction. Models were computed for three thresholds (5%, 10%, and 25%), with all resulting p-values adjusted for multiple comparisons using the FDR procedure (Benjamini and Hochberg, 1995). Only results for the 10% thresholds are shown, with 5% and 25% in Supplementary Figures.

## Acknowledgements

This work was supported by the European Research Council under the European Union’s Seventh Framework Programme (FP7/2007–2013)/ERC starting grant (grant number 716862) attributed to Mathilde Bonnefond and the Fondation pour la Recherche Medicale - grant ID FDT202106013010 awarded to Tommy Clausner. This work was conducted in the framework of the LabEx Cortex (“Construction, Function and Cognitive Function and Rehabilitation of the Cortex,” ANR-10-LABX-0042) of Université de Lyon. The authors would like to thank Jan Mathijs Schoffelen, Simon Homölle and Robert Oostenveld for their support with EEG source modelling. Furthermore, we would like to thank Rüdiger Stirnberg for his support on setting up the interleaved fMRI sequence, as well as Tim van Mourik for his support on cortical layer segmentation and Laurentius Huber for providing the crucial hint on accurate fMRI sub-millimetre within and between block volume (co-) registration and motion parameter estimation. Furthermore, the authors would like to thank Koen Haak for fruitful discussions and vital support in setting up the pRF recording pipeline, as well as Matthias Ekman for fruitful discussions on the improvement of the pRF mapping analysis pipeline. Lastly, the authors would like to thank Ole Jensen, Floris de Lange, David Norris and Jérémie Mattout for fruitful discussions and Paul Gaalman and the DCCN technical group for outstanding support.

## Supplementary Figures

**Figure 1:**
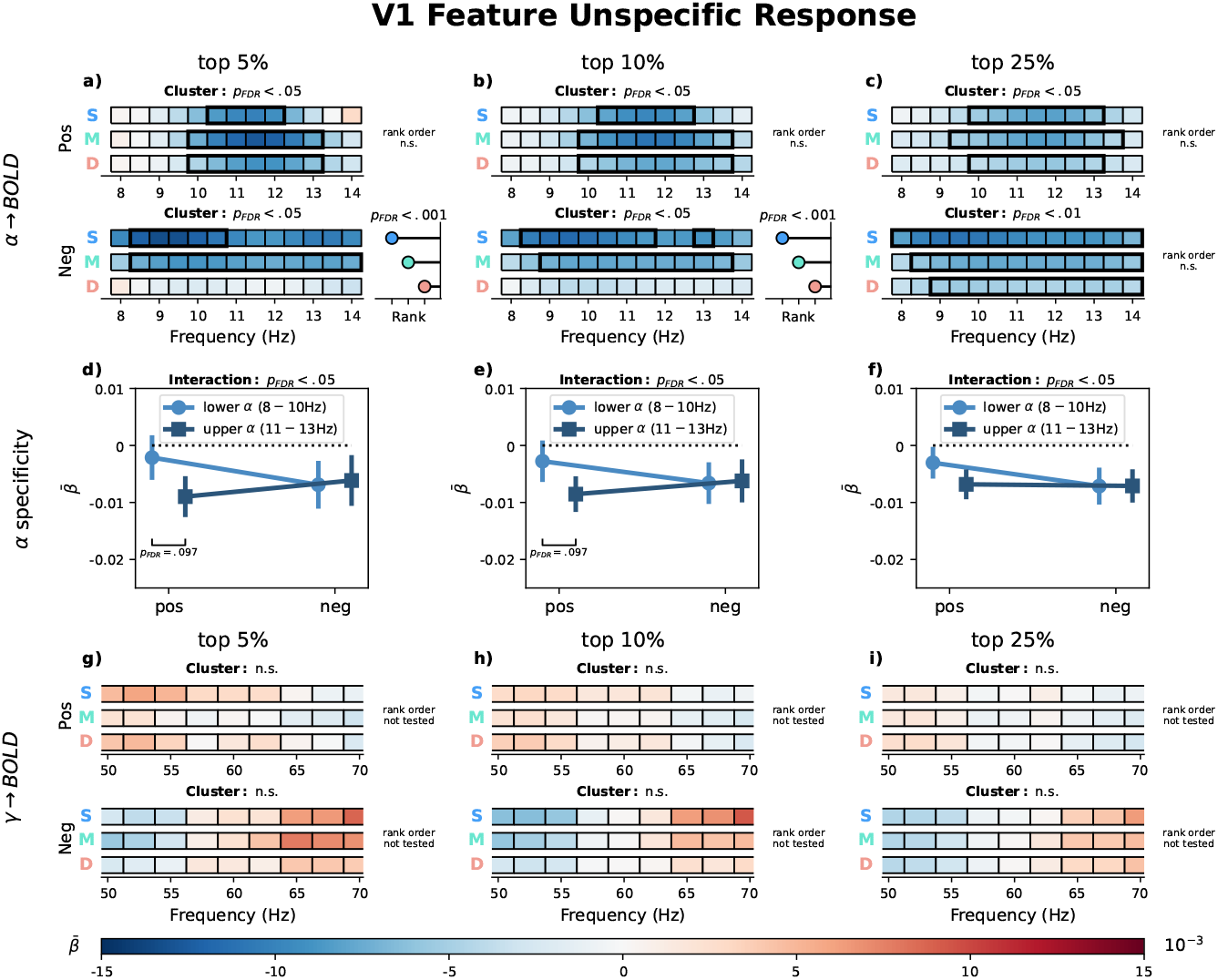
V1 feature-unspecific relationship between laminar BOLD signal and EEG. Average 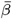 coefficients from a GLM predicting laminar BOLD signals in V1 from trial-by-trial EEG spectral power, shown separately for voxels with a strictly positive (Pos) or strictly negative (Neg) response to both stimulus orientations. Columns correspond to voxel-selection thresholds (top 5%, 10%, 25% most extreme t-values). 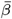 coefficients were weighted by laminar contribution weights, yielding estimates for superficial (**S**), middle (**M**), and deep (**D**) layers. **a–c**: *α*-band (8 – 14 Hz) layer *×* frequency plots. **d–f**: average 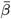 (*±* SEM) for lower (8 − 10 Hz) and upper (11 – 13 Hz) *α* sub-bands per voxel selection, tested with a linear mixed-effects model. **g–i**: *γ*-band (50 – 70 Hz) layer *×* frequency plots. Black rectangles indicate significant clusters (Maris and Oostenveld, 2007); where significant, lollipops to the right show the layer rank order of the effect (Clausner and Gentili, 2022). All p-values are FDR-corrected (Benjamini and Hochberg, 1995).

**Figure 2:**
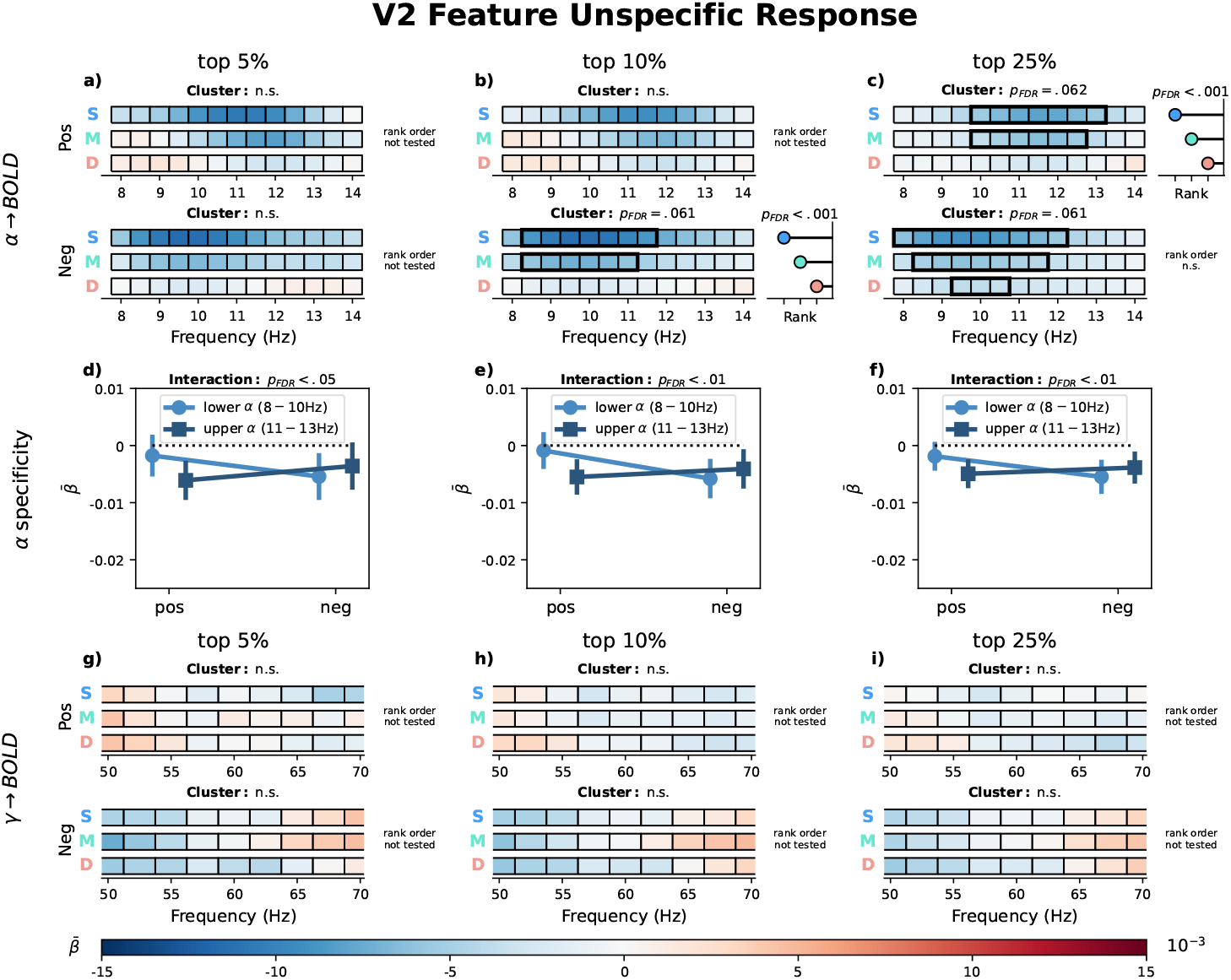
V2 feature-unspecific relationship between laminar BOLD signal and EEG. Average 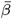 coefficients from a GLM predicting laminar BOLD signals in V2 from trial-by-trial EEG spectral power, shown separately for voxels with a strictly positive (Pos) or strictly negative (Neg) response to both stimulus orientations. Columns correspond to voxel-selection thresholds (top 5%, 10%, 25% most extreme t-values). 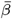 coefficients were weighted by laminar contribution weights, yielding estimates for superficial (**S**), middle (**M**), and deep (**D**) layers. **a–c**: *α*-band (8 – 14 Hz) layer × frequency plots. **d–f**: average 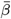 (*±* SEM) for lower (8 – 10 Hz) and upper (11 – 13 Hz) *α* sub-bands per voxel selection, tested with a linear mixed-effects model. **g–i**: *γ*-band (50 – 70 Hz) layer *×* frequency plots. Black rectangles indicate significant clusters (Maris and Oostenveld, 2007); where significant, lollipops to the right show the layer rank order of the effect (Clausner and Gentili, 2022). All p-values are FDR-corrected (Benjamini and Hochberg, 1995).

**Figure 3:**
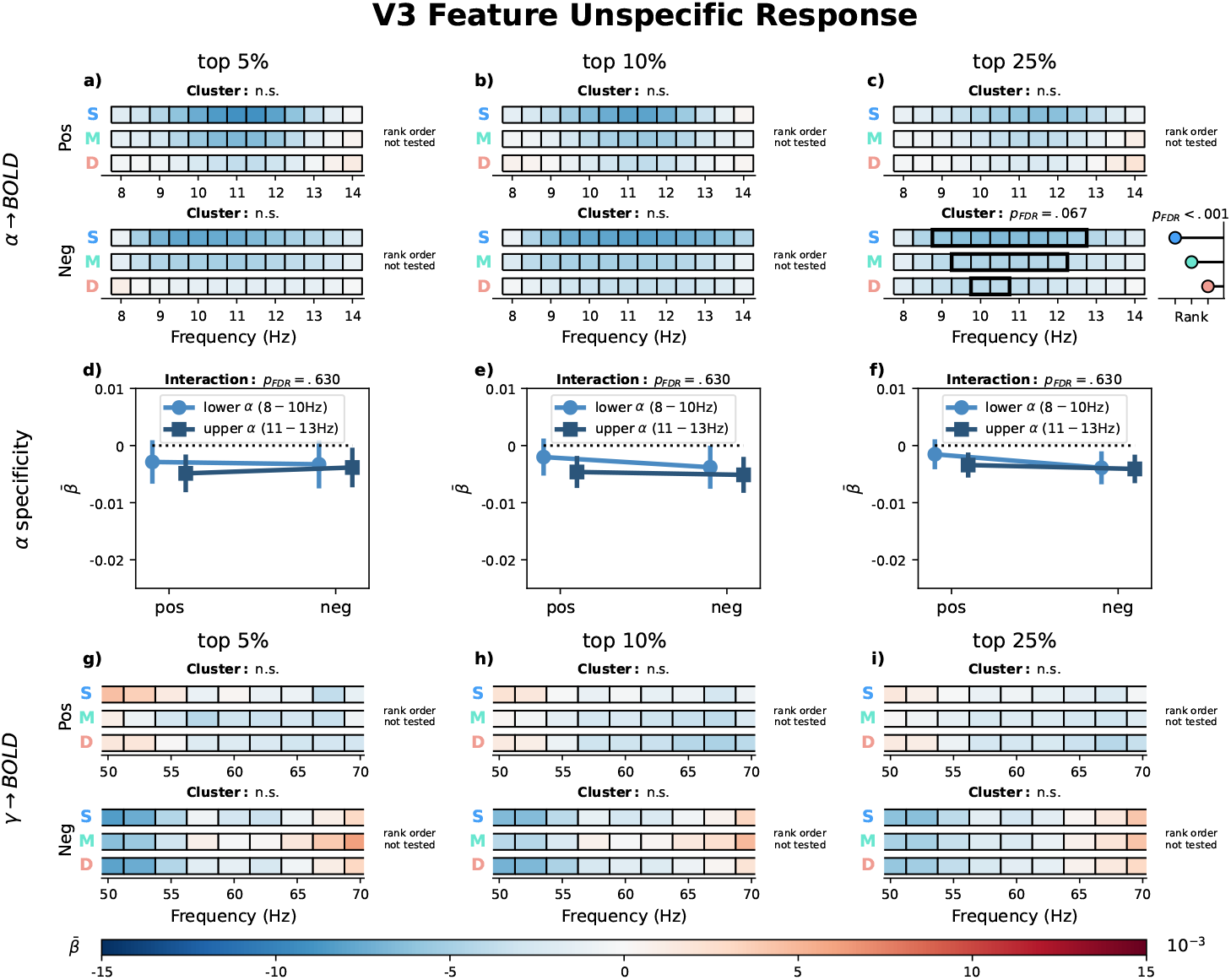
V3 feature-unspecific relationship between laminar BOLD signal and EEG. Average 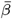 coefficients from a GLM predicting laminar BOLD signals in V3 from trial-by-trial EEG spectral power, shown separately for voxels with a strictly positive (Pos) or strictly negative (Neg) response to both stimulus orientations. Columns correspond to voxel-selection thresholds (top 5%, 10%, 25% most extreme t-values). 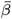 coefficients were weighted by laminar contribution weights, yielding estimates for superficial (**S**), middle (**M**), and deep (**D**) layers. **a–c**: *α*-band (8 – 14 Hz) layer × frequency plots. **d–f**: average 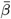 (*±* SEM) for lower (8 – 10 Hz) and upper (11 – 13 Hz) *α* sub-bands per voxel selection, tested with a linear mixed-effects model. **g–i**: *γ*-band (50 – 70 Hz) layer × frequency plots. Black rectangles indicate significant clusters (Maris and Oostenveld, 2007); where significant, lollipops to the right show the layer rank order of the effect (Clausner and Gentili, 2022). All p-values are FDR-corrected (Benjamini and Hochberg, 1995).

**Figure 4:**
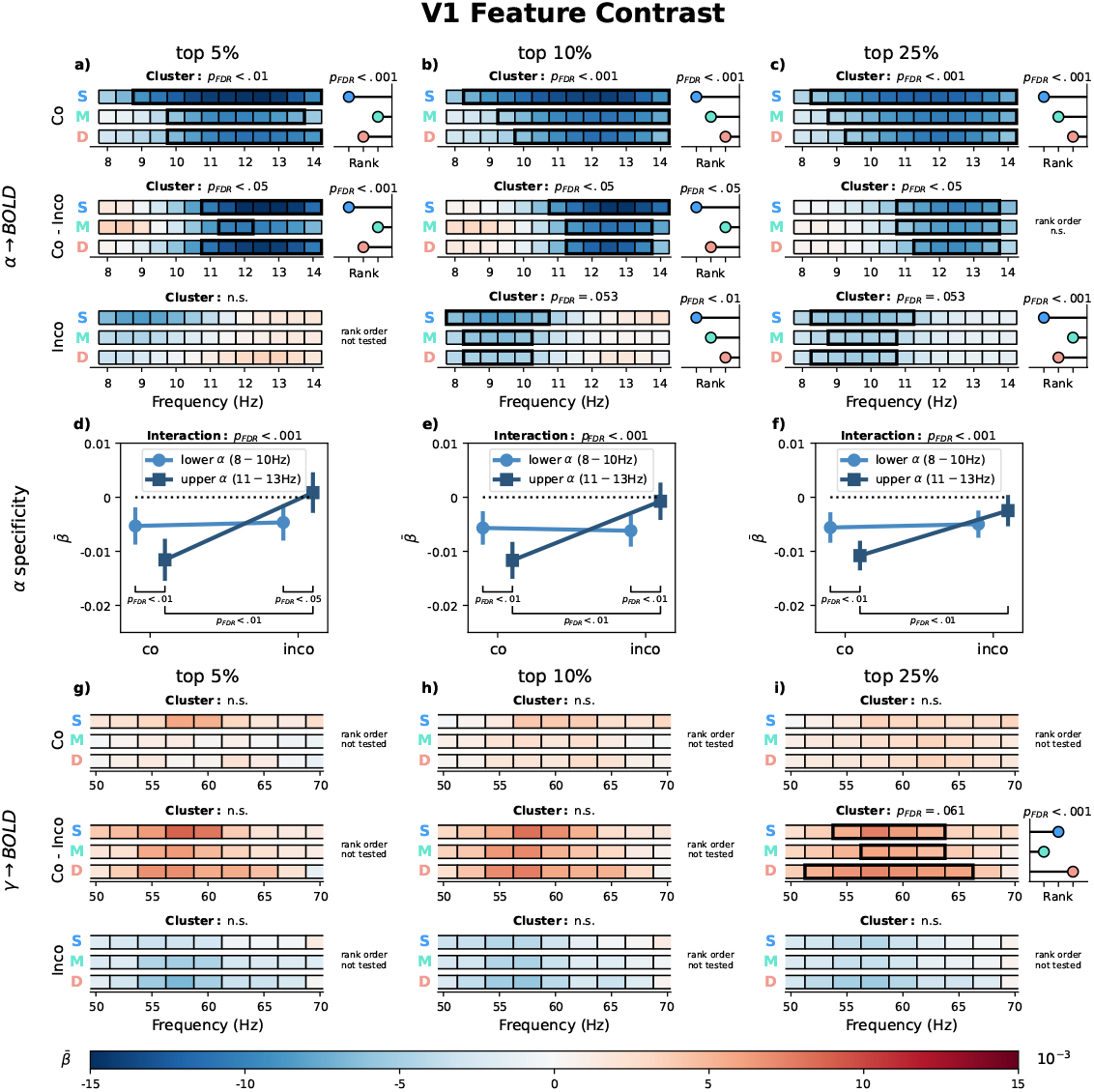
V1 relationship between laminar BOLD signal and EEG for the feature-specific contrast. Average 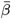 coefficients from a GLM predicting laminar BOLD signals in V1 from trial-by-trial EEG spectral power. Voxels where selected based on the first level fMRI contrast of *left* – *right* oriented stimuli (feature contrast). Columns correspond to voxel-selection thresholds (top 5%, 10%, 25% most extreme t-values). The GLM was computed using voxels with a stronger response to one orientation over the other. EEG regressors were computed from trials of the Congruent orientation (matching the preferred voxel orientation), the Incongruent orientation (opposite). Furthermore, the difference between congruent and incongruent models has been computed (Congruent - Incongruent). 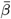 coefficients were weighted by laminar contribution weights, yielding estimates for superficial (**S**), middle (**M**), and deep (**D**) layers. **a–c**: *α*-band (8 – 14 Hz) layer × frequency plots. **d–f**: average 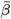 (*±* SEM) for lower (8 – 10 Hz) and upper (11 – 13 Hz) *α* sub-bands per voxel selection, tested with a linear mixed-effects model. **g–i**: *γ*-band (50 – 70 Hz) layer × frequency plots. Black rectangles indicate significant clusters (Maris and Oostenveld, 2007); where significant, lollipops to the right show the layer rank order of the effect (Clausner and Gentili, 2022). All p-values are FDR-corrected (Benjamini and Hochberg, 1995).

**Figure 5:**
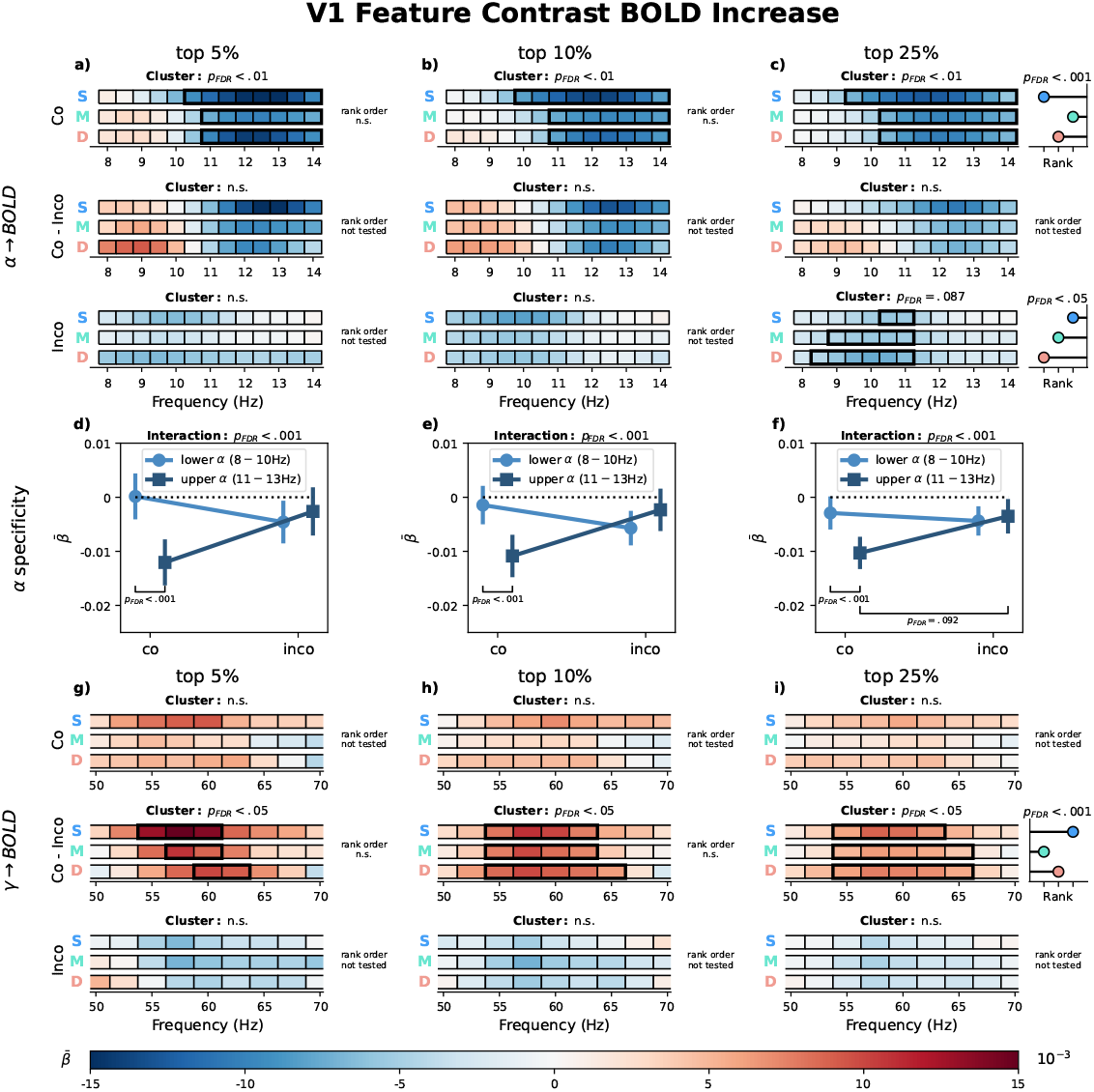
V1 relationship between laminar BOLD signal and EEG for the feature-specific contrast. Average 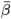 coefficients from a GLM predicting laminar BOLD signals in V1 from trial-by-trial EEG spectral power. Voxels where selected based on the first level fMRI contrast of *left* – *right* oriented stimuli and their (positive) response to either stimulus orientation (feature contrast BOLD increase). Columns correspond to voxel-selection thresholds (top 5%, 10%, 25% most extreme t-values). The GLM was computed using voxels with a stronger response to one orientation over the other. EEG regressors were computed from trials of the Congruent orientation (matching the preferred voxel orientation), the Incongruent orientation (opposite). Furthermore, the difference between congruent and incongruent models has been computed (Congruent - Incongruent). 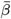 coefficients were weighted by laminar contribution weights, yielding estimates for superficial (**S**), middle (**M**), and deep (**D**) layers. **a–c**: *α*-band (8 – 14 Hz) layer × frequency plots. **d–f**: average 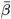 (*±* SEM) for lower (8 – 10 Hz) and upper (11 – 13 Hz) *α* sub-bands per voxel selection, tested with a linear mixed-effects model. **g–i**: *γ*-band (50 – 70 Hz) layer × frequency plots. Black rectangles indicate significant clusters (Maris and Oostenveld, 2007); where significant, lollipops to the right show the layer rank order of the effect (Clausner and Gentili, 2022). All p-values are FDR-corrected (Benjamini and Hochberg, 1995).

**Figure 6:**
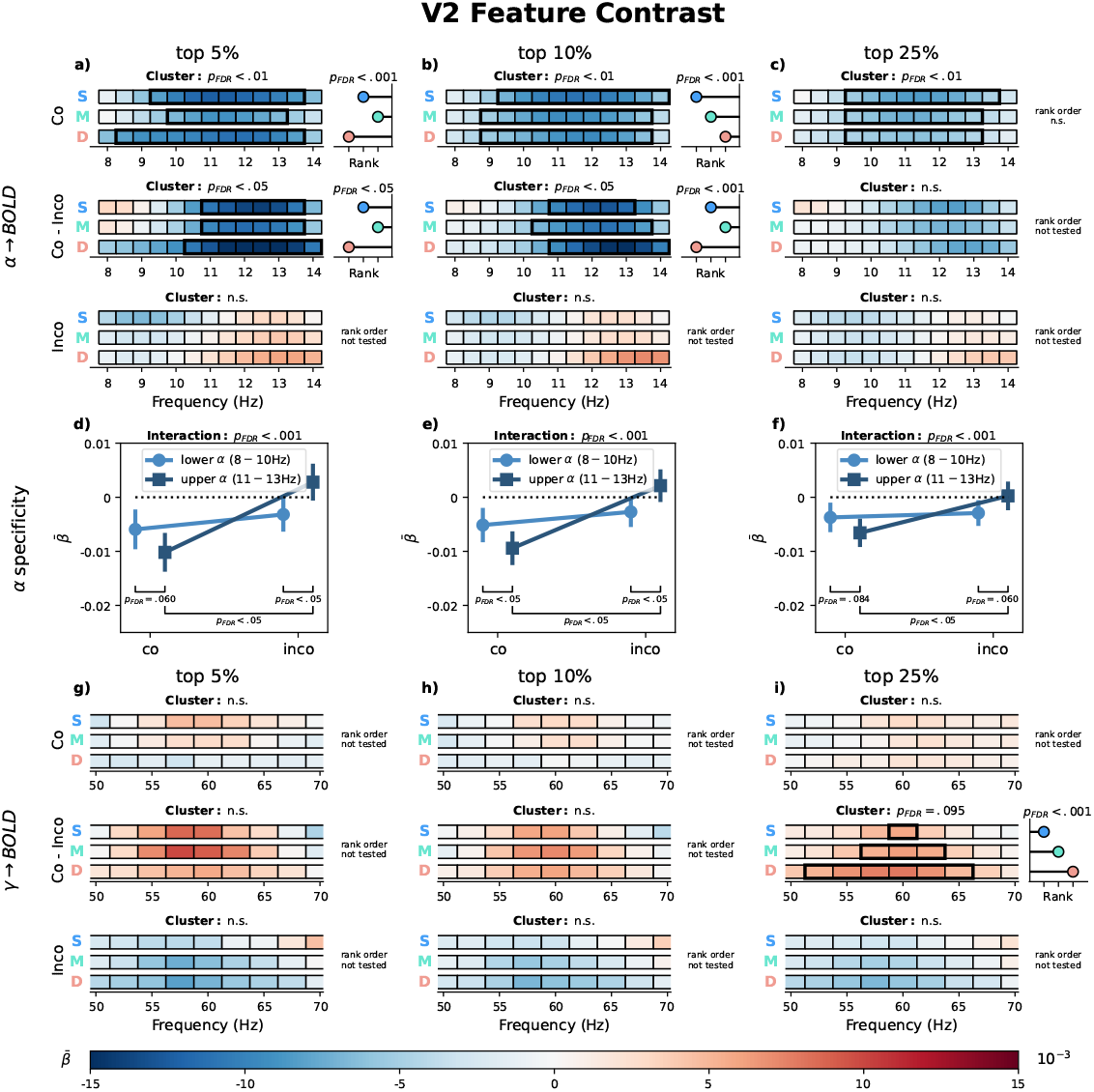
V2 relationship between laminar BOLD signal and EEG for the feature-specific contrast. Average 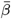 coefficients from a GLM predicting laminar BOLD signals in V2 from trial-by-trial EEG spectral power. Voxels where selected based on the first level fMRI contrast of *left* – *right* oriented stimuli (feature contrast). Columns correspond to voxel-selection thresholds (top 5%, 10%, 25% most extreme t-values). The GLM was computed using voxels with a stronger response to one orientation over the other. EEG regressors were computed from trials of the Congruent orientation (matching the preferred voxel orientation), the Incongruent orientation (opposite). Furthermore, the difference between congruent and incongruent models has been computed (Congruent - Incongruent). 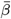 coefficients were weighted by laminar contribution weights, yielding estimates for superficial (**S**), middle (**M**), and deep (**D**) layers. **a–c**: *α*-band (8 – 14 Hz) layer × frequency plots. **d–f**: average 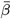 (*±* SEM) for lower (8 – 10 Hz) and upper (11 – 13 Hz) *α* sub-bands per voxel selection, tested with a linear mixed-effects model. **g–i**: *γ*-band (50 – 70 Hz) layer × frequency plots. Black rectangles indicate significant clusters (Maris and Oostenveld, 2007); where significant, lollipops to the right show the layer rank order of the effect (Clausner and Gentili, 2022). All p-values are FDR-corrected (Benjamini and Hochberg, 1995).

**Figure 7:**
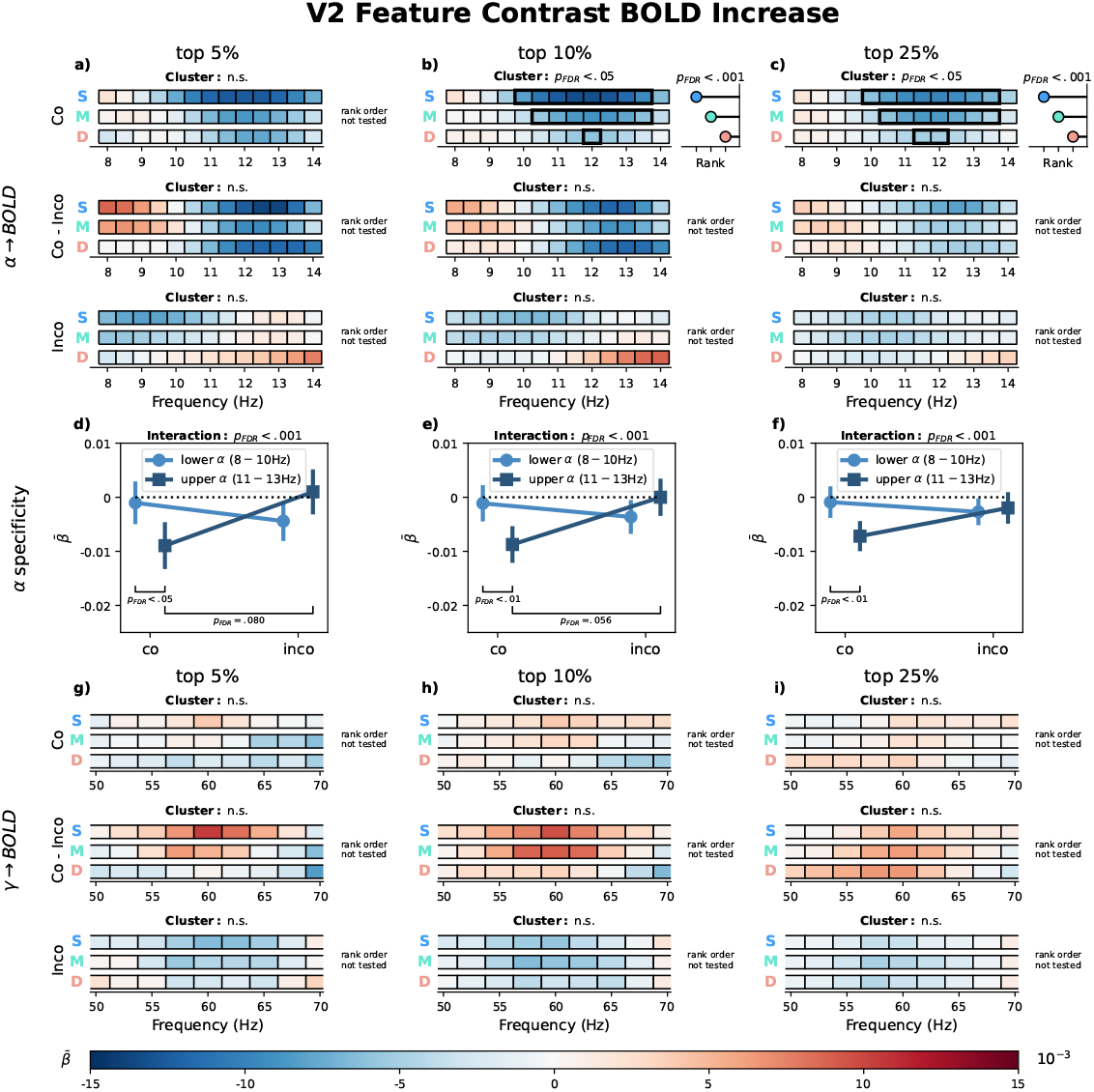
V2 relationship between laminar BOLD signal and EEG for the feature-specific contrast. Average 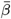 coefficients from a GLM predicting laminar BOLD signals in V2 from trial-by-trial EEG spectral power. Voxels where selected based on the first level fMRI contrast of *left* – *right* oriented stimuli and their (positive) response to either stimulus orientation (feature contrast BOLD increase). Columns correspond to voxel-selection thresholds (top 5%, 10%, 25% most extreme t-values). The GLM was computed using voxels with a stronger response to one orientation over the other. EEG regressors were computed from trials of the Congruent orientation (matching the preferred voxel orientation), the Incongruent orientation (opposite). Furthermore, the difference between congruent and incongruent models has been computed (Congruent - Incongruent). 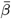 coefficients were weighted by laminar contribution weights, yielding estimates for superficial (**S**), middle (**M**), and deep (**D**) layers. **a–c**: *α*-band (8 – 14 Hz) layer × frequency plots. **d–f**: average 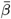 (*±* SEM) for lower (8 – 10 Hz) and upper (11 – 13 Hz) *α* sub-bands per voxel selection, tested with a linear mixed-effects model. **g–i**: *γ*-band (50 – 70 Hz) layer × frequency plots. Black rectangles indicate significant clusters (Maris and Oostenveld, 2007); where significant, lollipops to the right show the layer rank order of the effect (Clausner and Gentili, 2022). All p-values are FDR-corrected (Benjamini and Hochberg, 1995).

**Figure 8:**
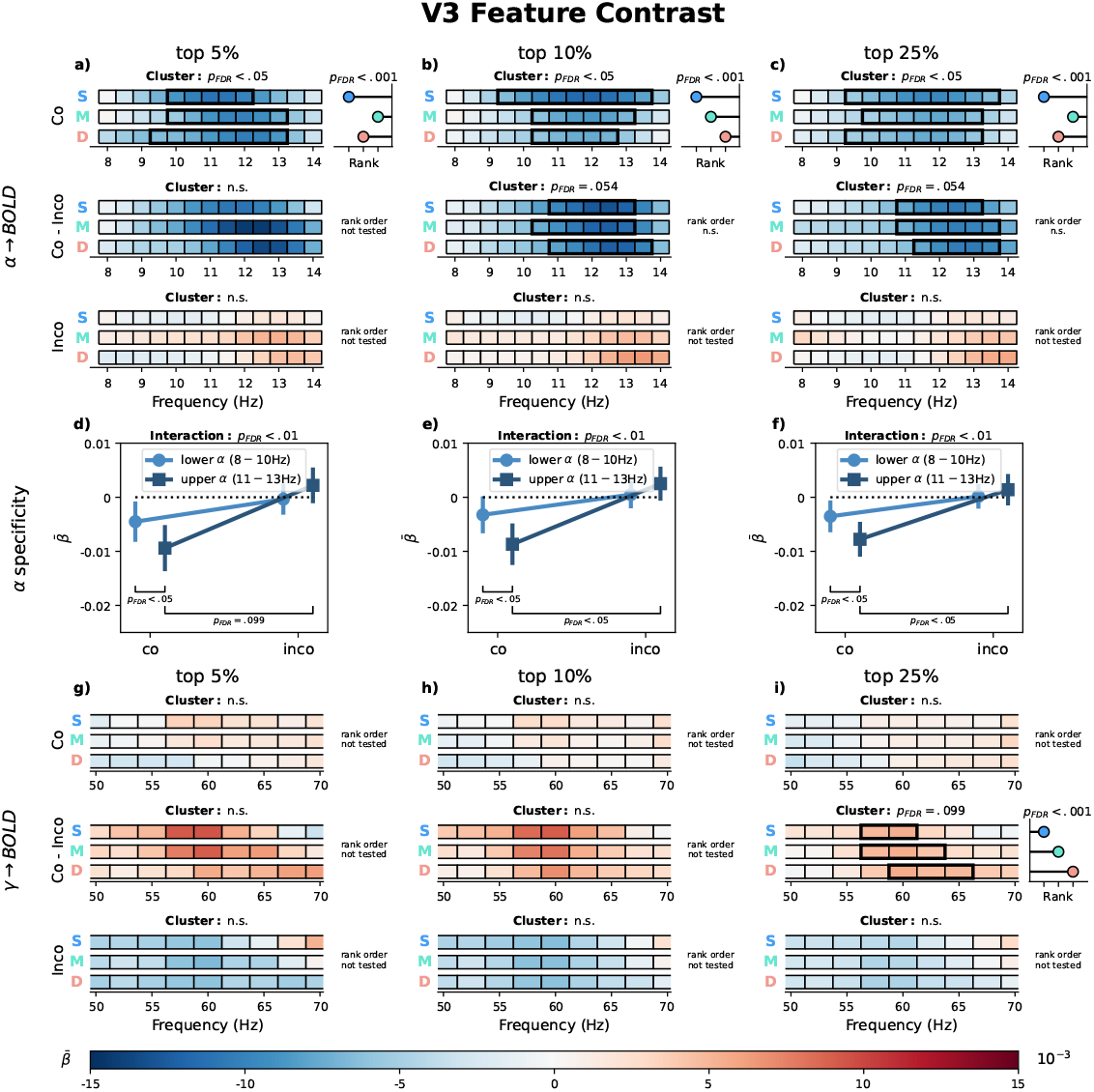
V3 relationship between laminar BOLD signal and EEG for the feature-specific contrast. Average 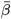 coefficients from a GLM predicting laminar BOLD signals in V3 from trial-by-trial EEG spectral power. Voxels where selected based on the first level fMRI contrast of *left* – *right* oriented stimuli (feature contrast). Columns correspond to voxel-selection thresholds (top 5%, 10%, 25% most extreme t-values). The GLM was computed using voxels with a stronger response to one orientation over the other. EEG regressors were computed from trials of the Congruent orientation (matching the preferred voxel orientation), the Incongruent orientation (opposite). Furthermore, the difference between congruent and incongruent models has been computed (Congruent - Incongruent). 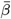 coefficients were weighted by laminar contribution weights, yielding estimates for superficial (**S**), middle (**M**), and deep (**D**) layers. **a–c**: *α*-band (8 – 14 Hz) layer × frequency plots. **d–f**: average 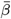 (*±* SEM) for lower (8 – 10 Hz) and upper (11 – 13 Hz) *α* sub-bands per voxel selection, tested with a linear mixed-effects model. **g–i**: *γ*-band (50 – 70 Hz) layer × frequency plots. Black rectangles indicate significant clusters (Maris and Oostenveld, 2007); where significant, lollipops to the right show the layer rank order of the effect (Clausner and Gentili, 2022). All p-values are FDR-corrected (Benjamini and Hochberg, 1995).

**Figure 9:**
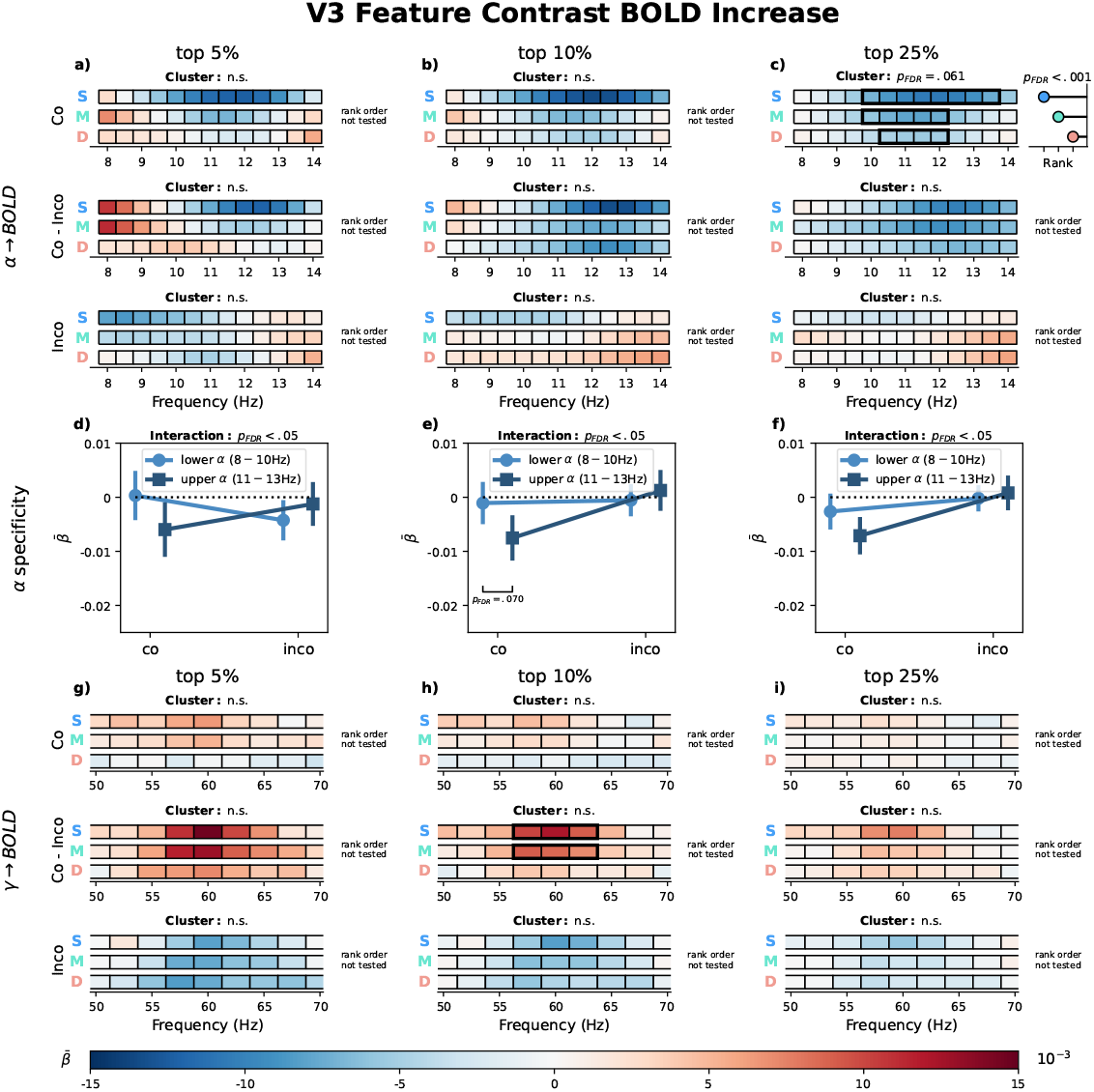
V3 relationship between laminar BOLD signal and EEG for the feature-specific contrast. Average 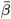 coefficients from a GLM predicting laminar BOLD signals in V3 from trial-by-trial EEG spectral power. Voxels where selected based on the first level fMRI contrast of *left right* oriented stimuli and their (positive) response to either stimulus orientation (feature contrast BOLD increase). Columns correspond to voxel-selection thresholds (top 5%, 10%, 25% most extreme t-values). The GLM was computed using voxels with a stronger response to one orientation over the other. EEG regressors were computed from trials of the Congruent orientation (matching the preferred voxel orientation), the Incongruent orientation (opposite). Furthermore, the difference between congruent and incongruent models has been computed (Congruent - Incongruent). 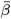 coefficients were weighted by laminar contribution weights, yielding estimates for superficial (**S**), middle (**M**), and deep (**D**) layers. **a–c**: *α*-band (8 – 14 Hz) layer × frequency plots. **d–f**: average 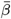 (*±* SEM) for lower (8 – 10 Hz) and upper (11 – 13 Hz) *α* sub-bands per voxel selection, tested with a linear mixed-effects model. **g–i**: *γ*-band (50 – 70 Hz) layer × frequency plots. Black rectangles indicate significant clusters (Maris and Oostenveld, 2007); where significant, lollipops to the right show the layer rank order of the effect (Clausner and Gentili, 2022). All p-values are FDR-corrected (Benjamini and Hochberg, 1995).

**Figure 10:**
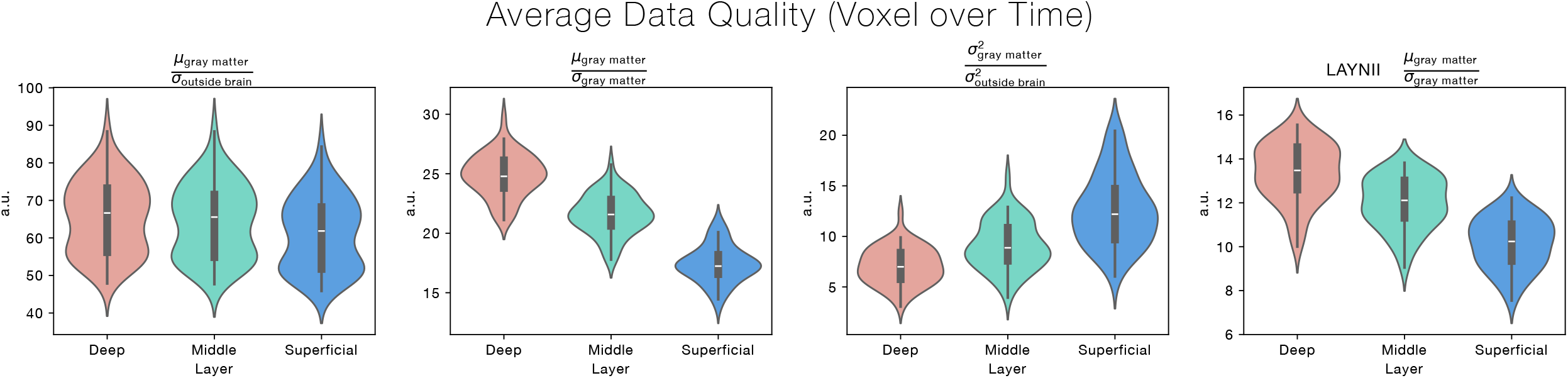
Signal-to-Noise Ratio. Several measures have been computed to estimate the temporal signal-to-noise ratio across layers, as indicated by the title of the respective sub-plot. All sub-plots except the rightmost sub-plot have been computed based on the used layering algorithm. We have included the tSNR estimate for the laminar segmentation as provided by LayNii (Huber et al., 2021), to allow for a comparison between our fraction-based approach and LayNii’s approach, which assigns entire voxels to specified layers. In general, the laminar profile of tSNR estimates was unexpected, because the vascular draining effect (Markuerkiaga et al., 2016) typically leads to higher signal changes in superficial layers. However, since both functionally relevant signal components, as well as physiological noise drains towards the cortical surface, a higher tSNR for deep layers is not entirely surprising. Importantly however, the tSNR profile is not reflected in our main results, which strengthens the overall validity.

**Figure 11:**
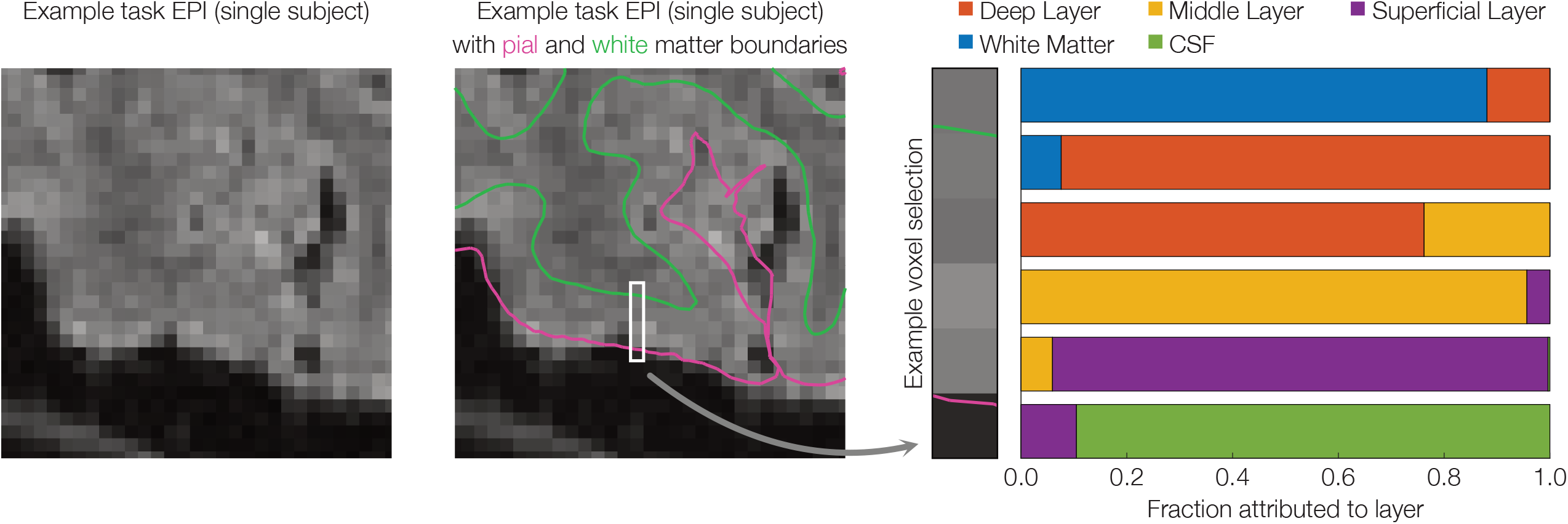
EPI example with layer segmentation. Left: a segment of a slice of a single participant’s raw fMRI volume during the task. Centre: The same segment, overlaid with pial and white matter boundaries that were used for reconstructing cortical depth. The white box indicates an example voxel selection for which the respective layer attributions are shown on the Right: Each voxel was assigned a fraction by how much this voxel belongs to a specific layer.

**Figure 12:**
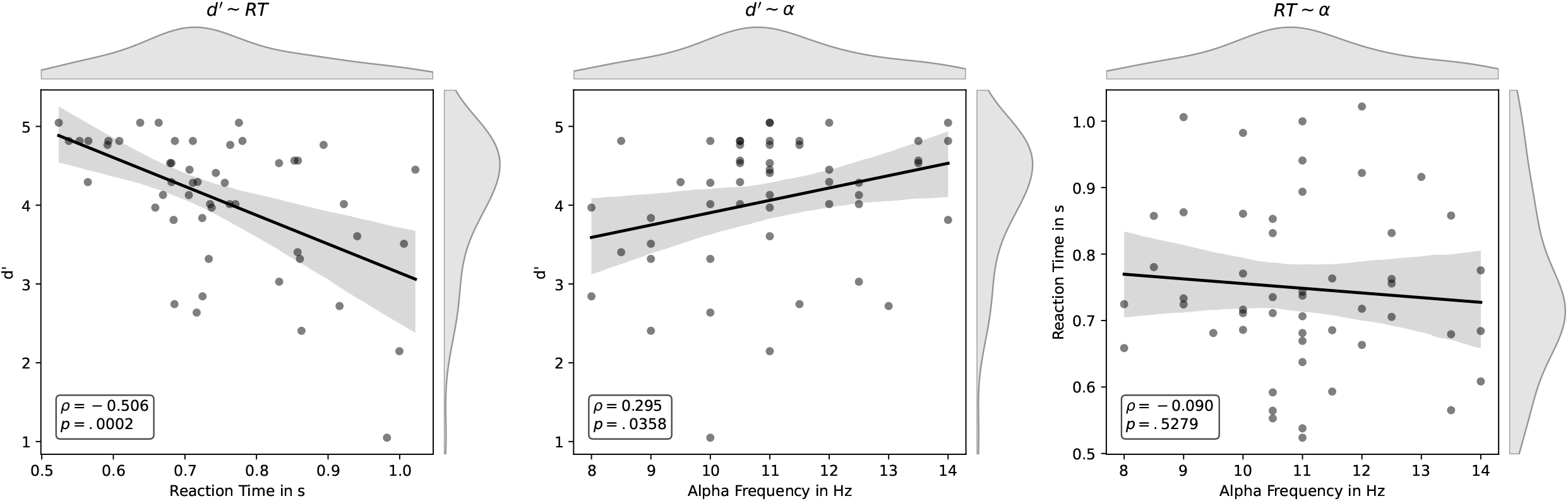
Correlation of *α* Frequency and Task Performance. Task performance for each participant has been defined as the average reaction time (RT) to correctly identified oddball stimuli, as well as d’ as a measure for average response accuracy. Individual *α* frequencies have been determined for each participant based on the average over all correctly identified non-oddball trials. Within the time of interest (0.1 to 0.8 s after stimulus onset) and frequencies of interest (8 to 14 Hz) the frequency with the largest decrease was chosen. Afterwards, per-participant frequencies have been correlated with per-participant average RTs and d’ values. We found that participants with the strongest decrease of *α* power in higher frequencies, also respond more accurately on average. No such correlation was observed between *α* frequency and RT. However, RT and d’ have been found to be negatively correlated as well.

**Figure 13:**
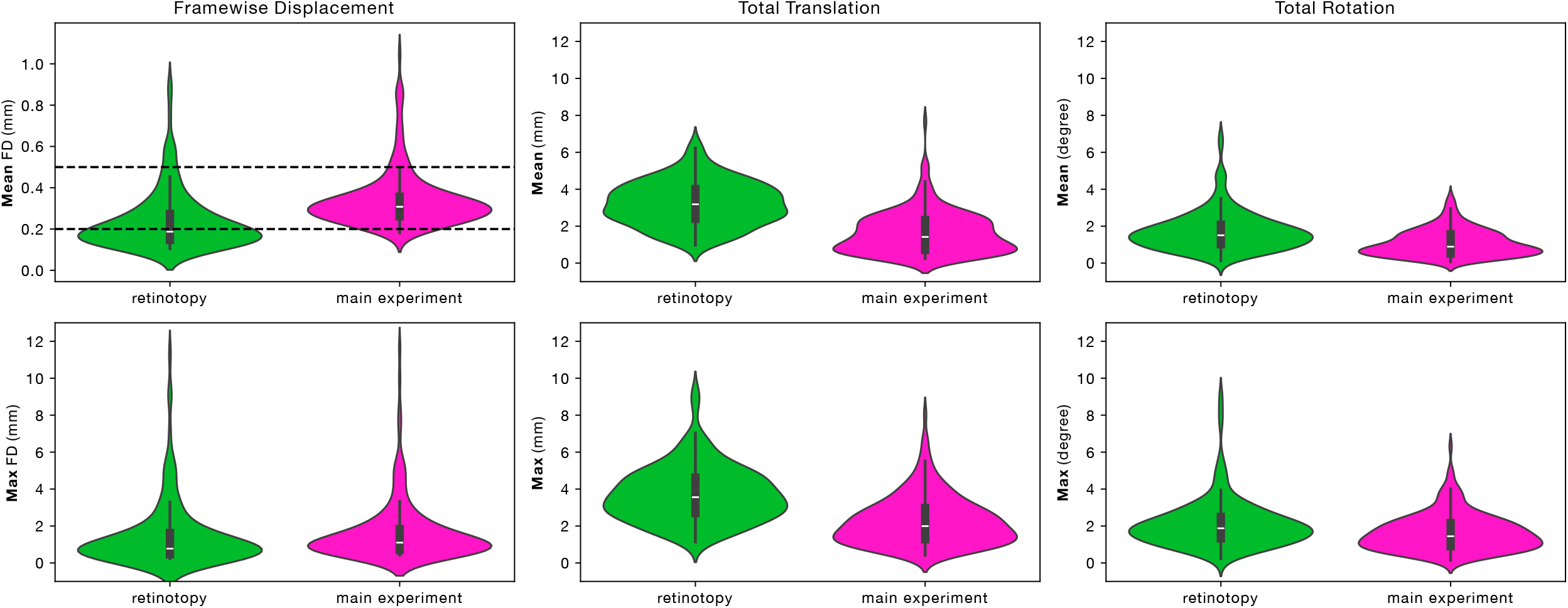
Average Motion of Participants. Average across participants for average and maximum frame-wise displacement (Power et al., 2012) using a radius of 50 mm, total translation and total rotation within one experimental block.

