## Supplementary Figures for "Frequency and Laminar Profile of Feature Specific Visual Activity Revealed by Interleaved EEG-fMRI"

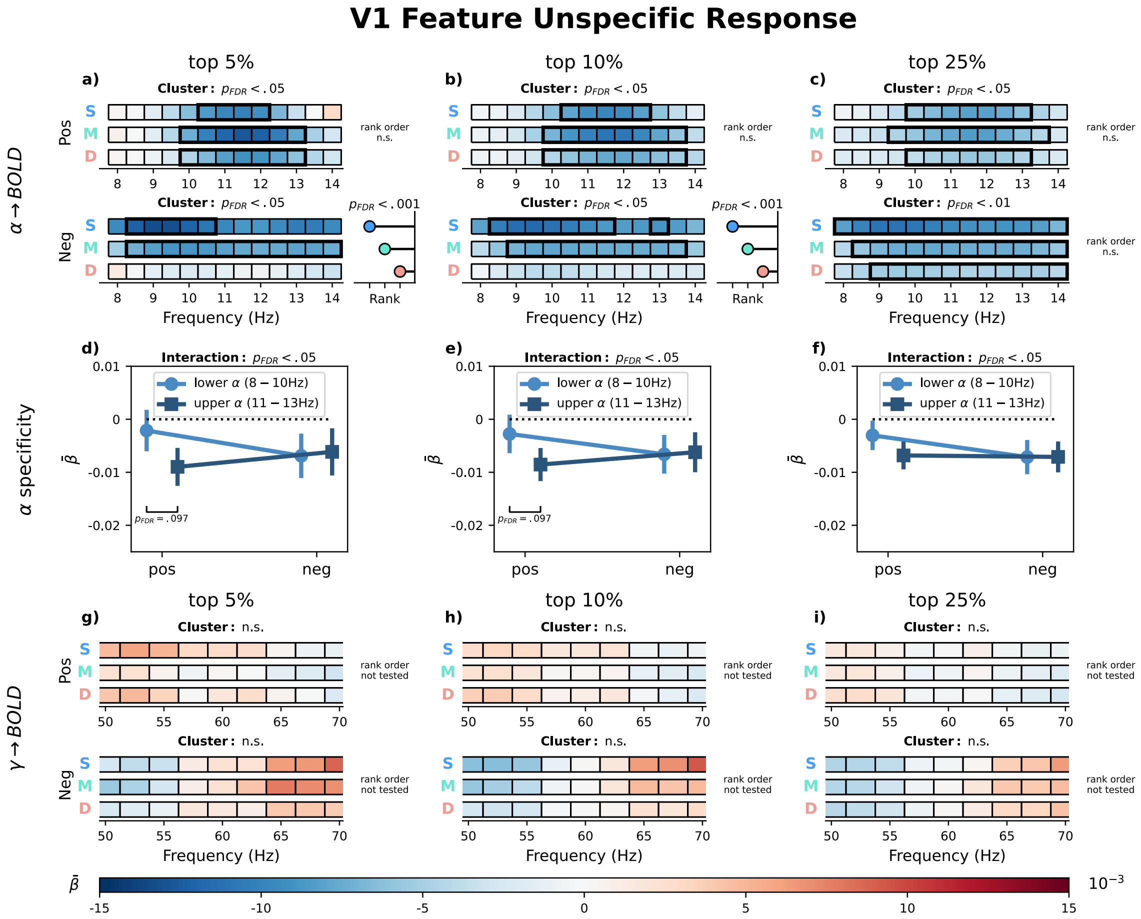


Figure 1: **V1 feature-unspecific relationship between laminar BOLD signal and EEG**. Average $\bar{\beta}$ coefficients from a GLM predicting laminar BOLD signals in V1 from trial-by-trial EEG spectral power, shown separately for voxels with a strictly positive (Pos) or strictly negative (Neg) response to both stimulus orientations. Columns correspond to voxel-selection thresholds (top 5%, 10%, 25% most extreme t-values). $\bar{\beta}$ coefficients were weighted by laminar contribution weights, yielding estimates for superficial (S), middle (M), and deep (D) layers. **a–c**: $\alpha$-band $(8 - 14\ \mathrm{Hz})$ layer × frequency plots. **d–f**: average $\bar{\beta}$ ($\pm$ SEM) for lower $(8 - 10\ \mathrm{Hz})$ and upper $(11 - 13\ \mathrm{Hz})$ $\alpha$ sub-bands per voxel selection, tested with a linear mixed-effects model. **g–i**: $\gamma$-band $(50 - 70\ \mathrm{Hz})$ layer × frequency plots. Black rectangles indicate significant clusters (Maris and Oostenveld, 2007); where significant, lollipops to the right show the layer rank order of the effect (Clausner and Gentili, 2022). All p-values are FDR-corrected (Benjamini and Hochberg, 1995).

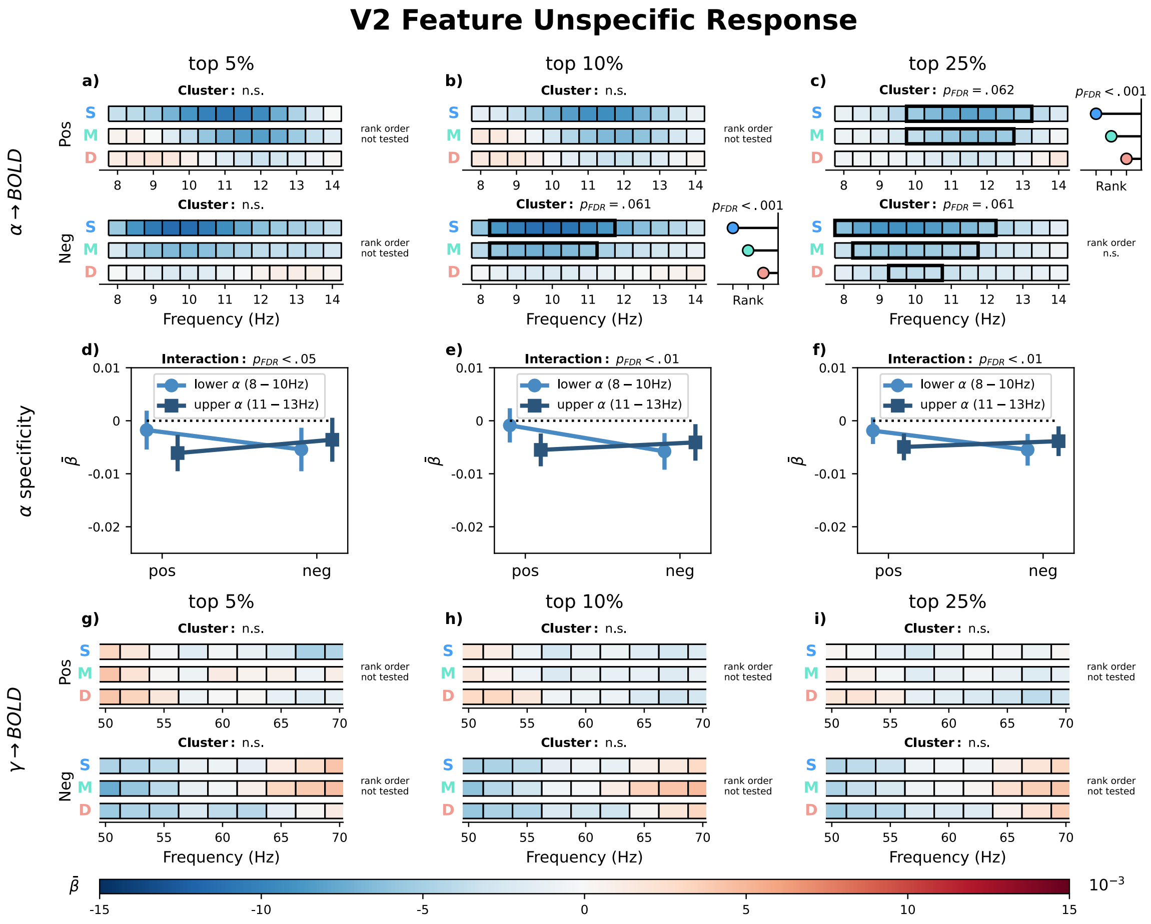


Figure 2: **V2 feature-unspecific relationship between laminar BOLD signal and EEG**. Average $\bar{\beta}$ coefficients from a GLM predicting laminar BOLD signals in V2 from trial-by-trial EEG spectral power, shown separately for voxels with a strictly positive (Pos) or strictly negative (Neg) response to both stimulus orientations. Columns correspond to voxel-selection thresholds (top 5%, 10%, 25% most extreme t-values). $\bar{\beta}$ coefficients were weighted by laminar contribution weights, yielding estimates for superficial (S), middle (M), and deep (D) layers. **a–c**: $\alpha$-band $(8 - 14\ \mathrm{Hz})$ layer × frequency plots. **d–f**: average $\bar{\beta}$ ($\pm$ SEM) for lower $(8 - 10\ \mathrm{Hz})$ and upper $(11 - 13\ \mathrm{Hz})$ $\alpha$ sub-bands per voxel selection, tested with a linear mixed-effects model. **g–i**: $\gamma$-band $(50 - 70\ \mathrm{Hz})$ layer × frequency plots. Black rectangles indicate significant clusters (Maris and Oostenveld, 2007); where significant, lollipops to the right show the layer rank order of the effect (Clausner and Gentili, 2022). All p-values are FDR-corrected (Benjamini and Hochberg, 1995).

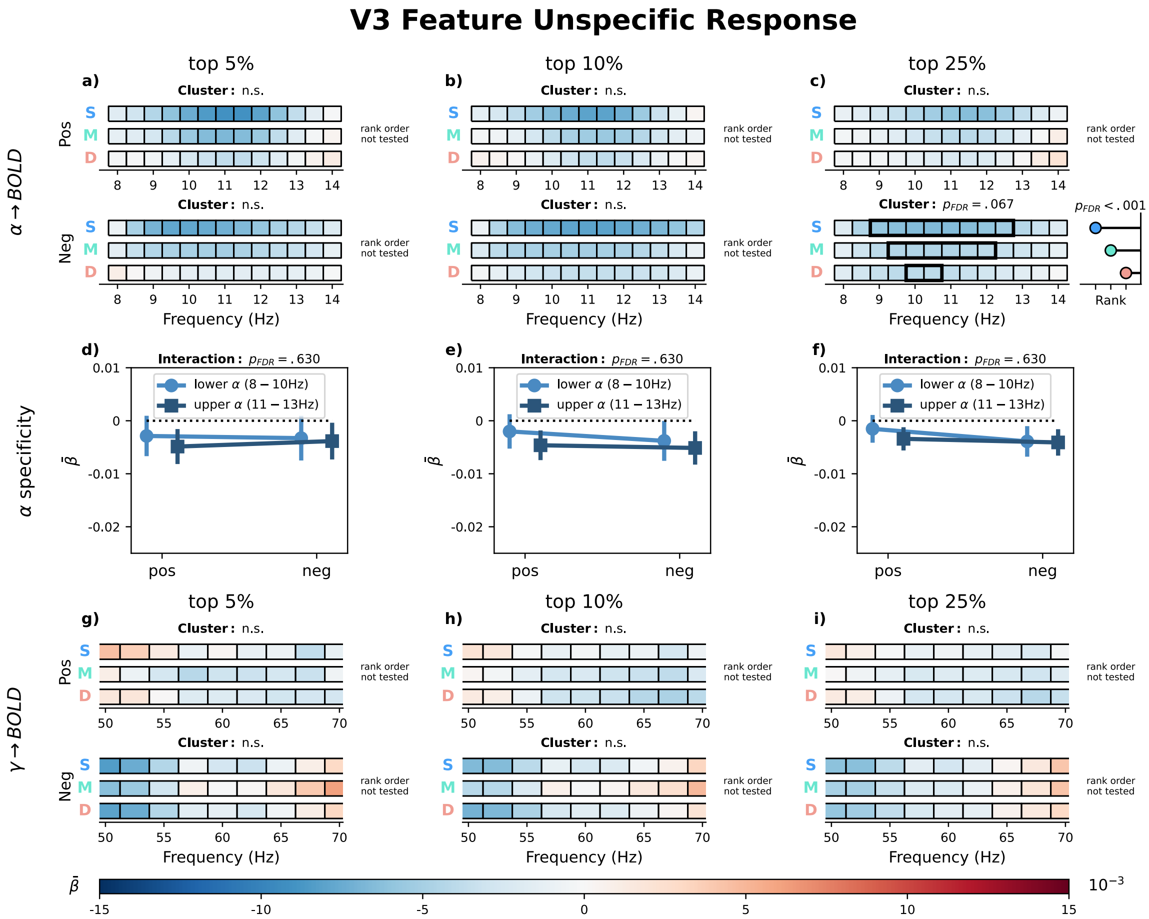


Figure 3: **V3 feature-unspecific relationship between laminar BOLD signal and EEG**. Average $\bar{\beta}$ coefficients from a GLM predicting laminar BOLD signals in V3 from trial-by-trial EEG spectral power, shown separately for voxels with a strictly positive (Pos) or strictly negative (Neg) response to both stimulus orientations. Columns correspond to voxel-selection thresholds (top 5%, 10%, 25% most extreme t-values). $\bar{\beta}$ coefficients were weighted by laminar contribution weights, yielding estimates for superficial (S), middle (M), and deep (D) layers. **a–c**: $\alpha$-band $(8 - 14\ \mathrm{Hz})$ layer × frequency plots. **d–f**: average $\bar{\beta}$ ($\pm$ SEM) for lower $(8 - 10\ \mathrm{Hz})$ and upper $(11 - 13\ \mathrm{Hz})$ $\alpha$ sub-bands per voxel selection, tested with a linear mixed-effects model. **g–i**: $\gamma$-band $(50 - 70\ \mathrm{Hz})$ layer × frequency plots. Black rectangles indicate significant clusters (Maris and Oostenveld, 2007); where significant, lollipops to the right show the layer rank order of the effect (Clausner and Gentili, 2022). All p-values are FDR-corrected (Benjamini and Hochberg, 1995).

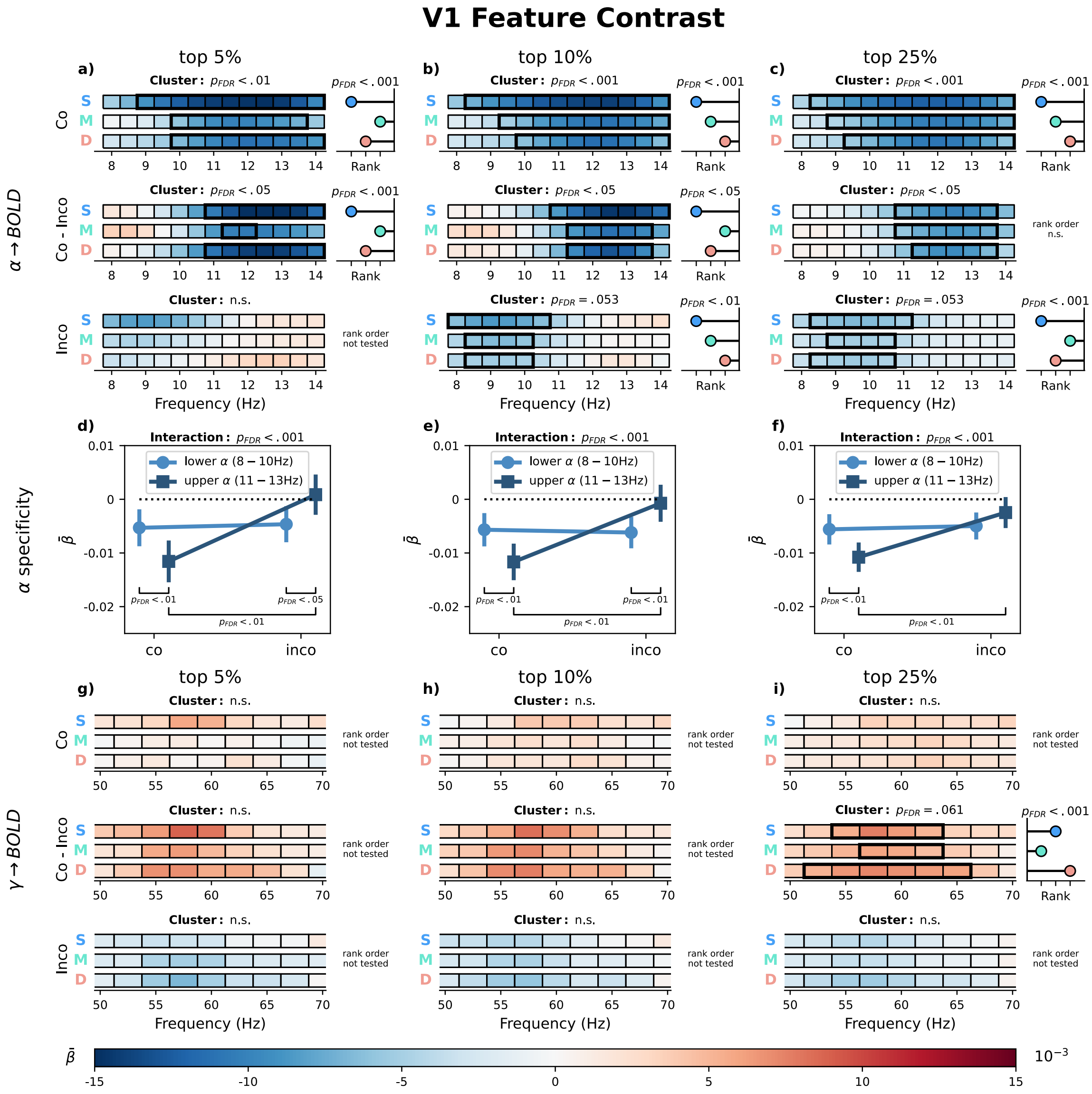


Figure 4: **V1 relationship between laminar BOLD signal and EEG for the feature-specific contrast**. Average $\bar{\beta}$ coefficients from a GLM predicting laminar BOLD signals in V1 from trial-by-trial EEG spectral power. Voxels where selected based on the first level fMRI contrast of $left - right$ oriented stimuli (feature contrast). Columns correspond to voxel-selection thresholds (top 5%, 10%, 25% most extreme t-values). The GLM was computed using voxels with a stronger response to one orientation over the other. EEG regressors were computed from trials of the Congruent orientation (matching the preferred voxel orientation), the Incongruent orientation (opposite). Furthermore, the difference between congruent and incongruent models has been computed (Congruent - Incongruent). $\bar{\beta}$ coefficients were weighted by laminar contribution weights, yielding estimates for superficial (**S**), middle (**M**), and deep (**D**) layers. **a–c**: $\alpha$-band $(8 - 14\ \mathrm{Hz})$ layer $\times$ frequency plots. **d–f**: average $\bar{\beta}$ ($\pm$ SEM) for lower $(8 - 10\ \mathrm{Hz})$ and upper $(11 - 13\ \mathrm{Hz})$ $\alpha$ sub-bands per voxel selection, tested with a linear mixed-effects model. **g–i**: $\gamma$-band $(50 - 70\ \mathrm{Hz})$ layer $\times$ frequency plots. Black rectangles indicate significant clusters (Maris and Oostenveld, 2007); where significant, lollipops to the right show the layer rank order of the effect (Clausner and Gentili, 2022). All p-values are FDR-corrected (Benjamini and Hochberg, 1995).

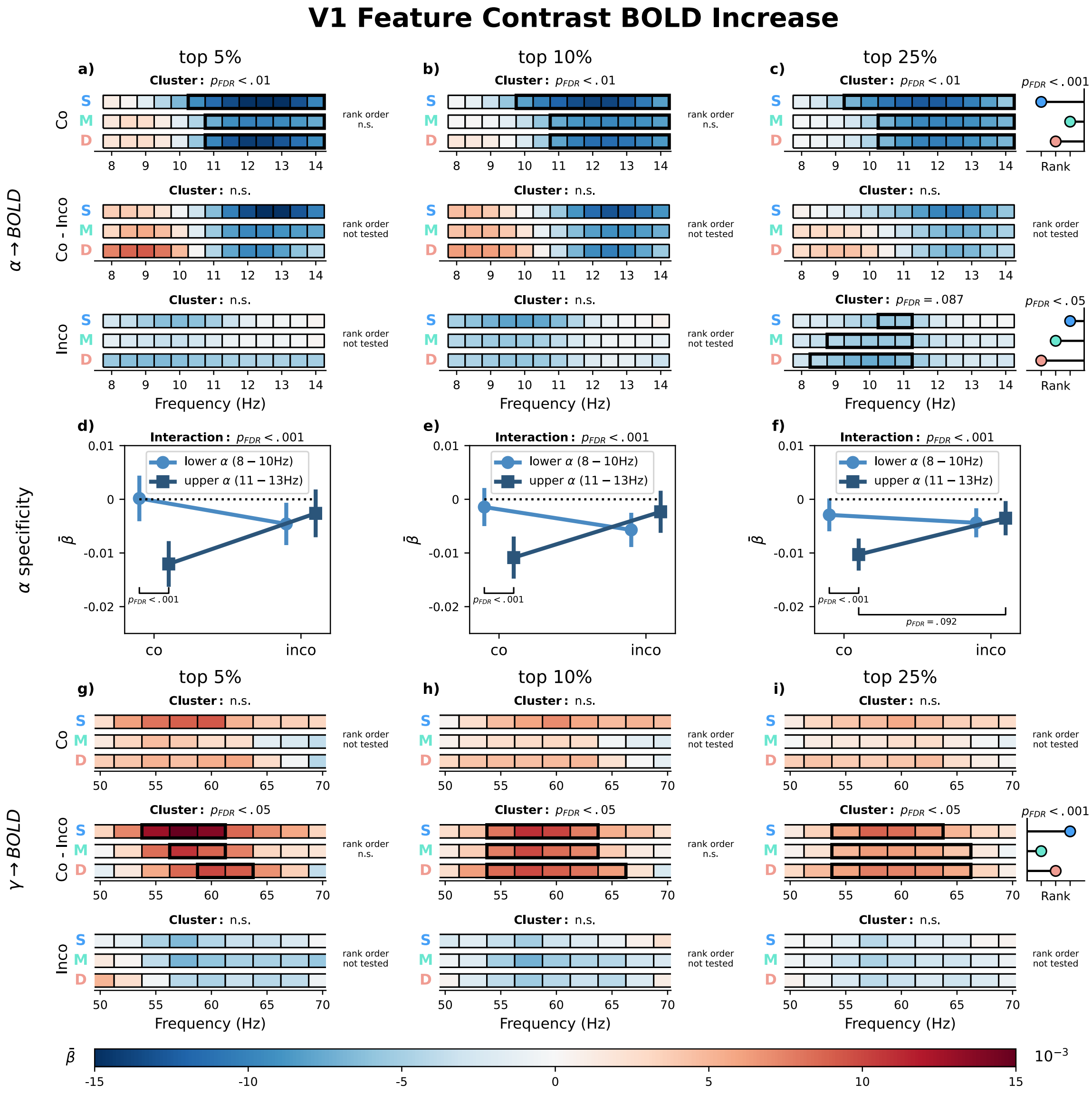


Figure 5: **V1 relationship between laminar BOLD signal and EEG for the feature-specific contrast**. Average $\bar{\beta}$ coefficients from a GLM predicting laminar BOLD signals in V1 from trial-by-trial EEG spectral power. Voxels where selected based on the first level fMRI contrast of $left - right$ oriented stimuli and their (positive) response to either stimulus orientation (feature contrast BOLD increase). Columns correspond to voxel-selection thresholds (top 5%, 10%, 25% most extreme t-values). The GLM was computed using voxels with a stronger response to one orientation over the other. EEG regressors were computed from trials of the Congruent orientation (matching the preferred voxel orientation), the Incongruent orientation (opposite). Furthermore, the difference between congruent and incongruent models has been computed (Congruent - Incongruent). $\bar{\beta}$ coefficients were weighted by laminar contribution weights, yielding estimates for superficial (S), middle (M), and deep (D) layers. **a–c**: $\alpha$-band $(8 - 14 \text{ Hz})$ layer $\times$ frequency plots. **d–f**: average $\bar{\beta}$ ($\pm$ SEM) for lower $(8 - 10 \text{ Hz})$ and upper $(11 - 13 \text{ Hz})$ $\alpha$ sub-bands per voxel selection, tested with a linear mixed-effects model. **g–i**: $\gamma$-band $(50 - 70 \text{ Hz})$ layer $\times$ frequency plots. Black rectangles indicate significant clusters (Maris and Oostenveld 2007); where significant, lollipops to the right show the layer rank order of the effect (Clausner and Gentili, 2022). All p-values are FDR-corrected (Benjamini and Hochberg, 1995).

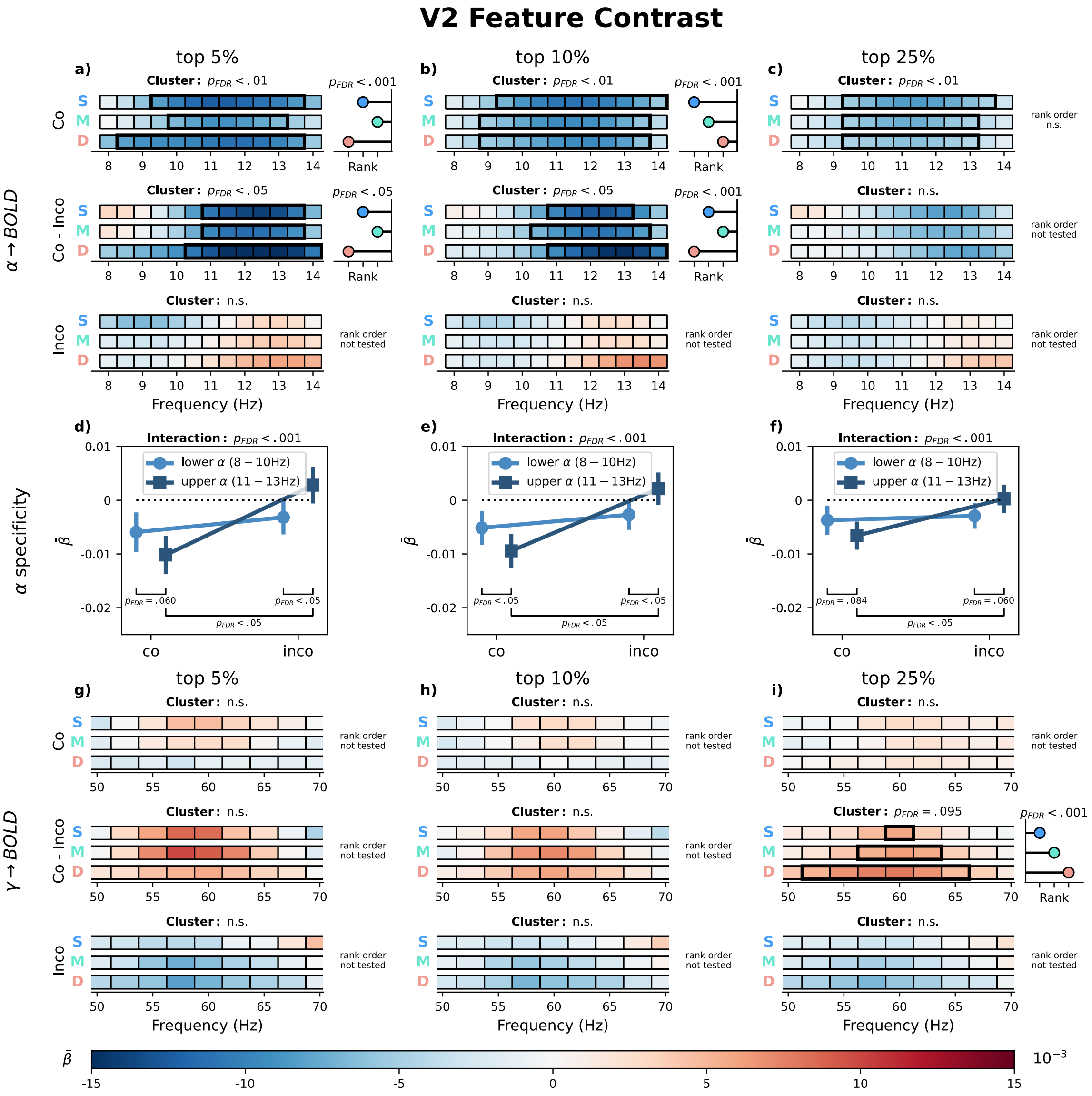


Figure 6: **V2 relationship between laminar BOLD signal and EEG for the feature-specific contrast**. Average $\bar{\beta}$ coefficients from a GLM predicting laminar BOLD signals in V2 from trial-by-trial EEG spectral power. Voxels where selected based on the first level fMRI contrast of $left - right$ oriented stimuli (feature contrast). Columns correspond to voxel-selection thresholds (top 5%, 10%, 25% most extreme t-values). The GLM was computed using voxels with a stronger response to one orientation over the other. EEG regressors were computed from trials of the Congruent orientation (matching the preferred voxel orientation), the Incongruent orientation (opposite). Furthermore, the difference between congruent and incongruent models has been computed (Congruent - Incongruent). $\bar{\beta}$ coefficients were weighted by laminar contribution weights, yielding estimates for superficial (**S**), middle (**M**), and deep (**D**) layers. **a–c**: $\alpha$-band $(8 - 14\ \mathrm{Hz})$ layer $\times$ frequency plots. **d–f**: average $\bar{\beta}$ ($\pm$ SEM) for lower $(8 - 10\ \mathrm{Hz})$ and upper $(11 - 13\ \mathrm{Hz})$ $\alpha$ sub-bands per voxel selection, tested with a linear mixed-effects model. **g–i**: $\gamma$-band $(50 - 70\ \mathrm{Hz})$ layer $\times$ frequency plots. Black rectangles indicate significant clusters (Maris and Oostenveld, 2007); where significant, lollipops to the right show the layer rank order of the effect (Clausner and Gentili, 2022). All p-values are FDR-corrected (Benjamini and Hochberg, 1995).

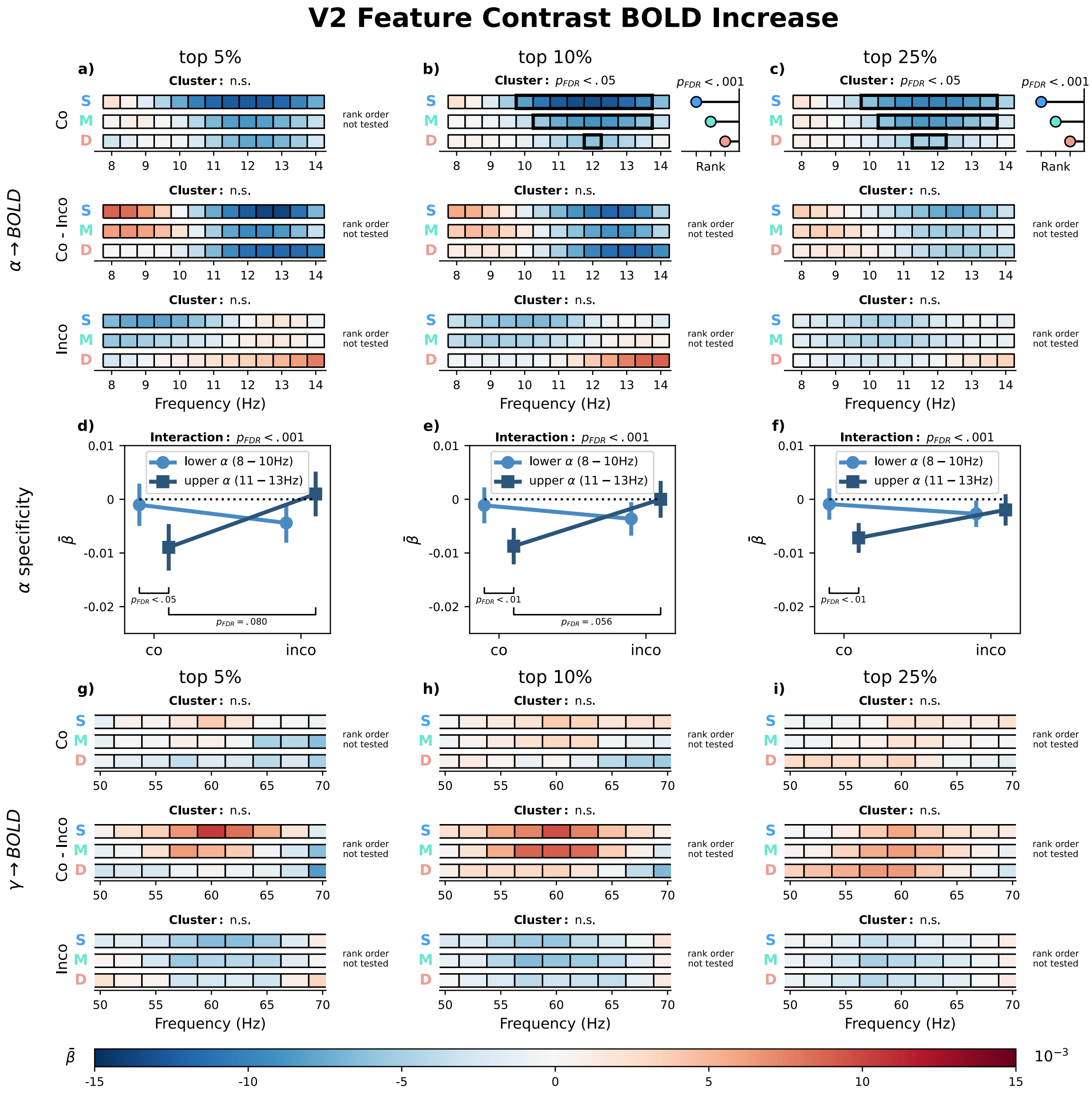


Figure 7: **V2 relationship between laminar BOLD signal and EEG for the feature-specific contrast**. Average $\bar{\beta}$ coefficients from a GLM predicting laminar BOLD signals in V2 from trial-by-trial EEG spectral power. Voxels where selected based on the first level fMRI contrast of $left - right$ oriented stimuli and their (positive) response to either stimulus orientation (feature contrast BOLD increase). Columns correspond to voxel-selection thresholds (top 5%, 10%, 25% most extreme t-values). The GLM was computed using voxels with a stronger response to one orientation over the other. EEG regressors were computed from trials of the Congruent orientation (matching the preferred voxel orientation), the Incongruent orientation (opposite). Furthermore, the difference between congruent and incongruent models has been computed (Congruent - Incongruent). $\bar{\beta}$ coefficients were weighted by laminar contribution weights, yielding estimates for superficial (S), middle (M), and deep (D) layers. **a–c**: $\alpha$-band ($8 - 14$ Hz) layer $\times$ frequency plots. **d–f**: average $\bar{\beta}$ ($\pm$ SEM) for lower ($8 - 10$ Hz) and upper ($11 - 13$ Hz) $\alpha$ sub-bands per voxel selection, tested with a linear mixed-effects model. **g–i**: $\gamma$-band ($50 - 70$ Hz) layer $\times$ frequency plots. Black rectangles indicate significant clusters (Maris and Oostenveld 2007); where significant, lollipops to the right show the layer rank order of the effect (Clausner and Gentili, 2022). All p-values are FDR-corrected (Benjamini and Hochberg, 1995).

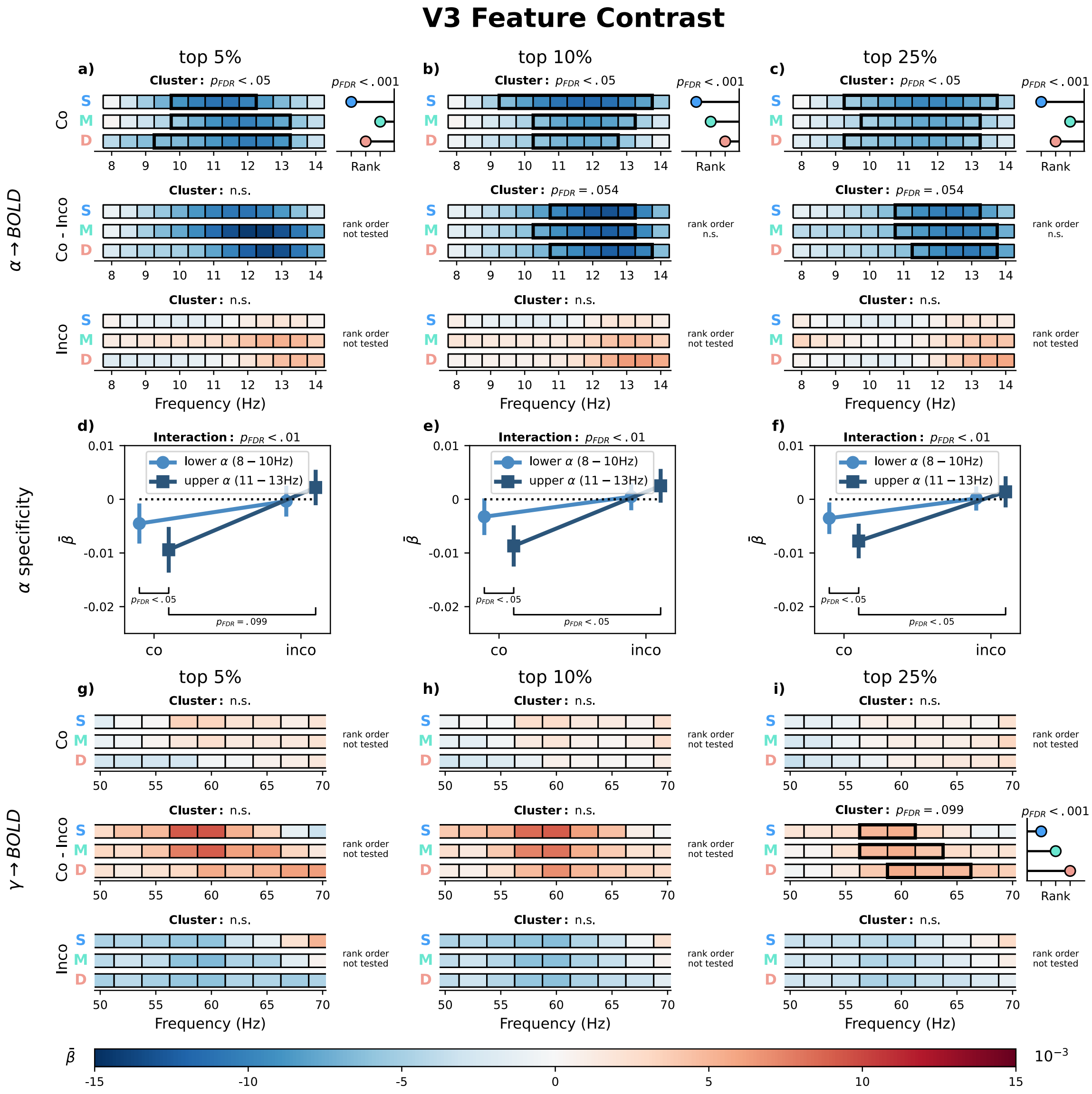


Figure 8: **V3 relationship between laminar BOLD signal and EEG for the feature-specific contrast**. Average $\bar{\beta}$ coefficients from a GLM predicting laminar BOLD signals in V3 from trial-by-trial EEG spectral power. Voxels where selected based on the first level fMRI contrast of $left - right$ oriented stimuli (feature contrast). Columns correspond to voxel-selection thresholds (top 5%, 10%, 25% most extreme t-values). The GLM was computed using voxels with a stronger response to one orientation over the other. EEG regressors were computed from trials of the Congruent orientation (matching the preferred voxel orientation), the Incongruent orientation (opposite). Furthermore, the difference between congruent and incongruent models has been computed (Congruent - Incongruent). $\bar{\beta}$ coefficients were weighted by laminar contribution weights, yielding estimates for superficial (S), middle (M), and deep (D) layers. **a–c**: $\alpha$-band $(8 - 14\ \mathrm{Hz})$ layer × frequency plots. **d–f**: average $\bar{\beta}$ (± SEM) for lower $(8 - 10\ \mathrm{Hz})$ and upper $(11 - 13\ \mathrm{Hz})$ $\alpha$ sub-bands per voxel selection, tested with a linear mixed-effects model. **g–i**: $\gamma$-band $(50 - 70\ \mathrm{Hz})$ layer × frequency plots. Black rectangles indicate significant clusters (Maris and Oostenveld, 2007); where significant, lollipops to the right show the layer rank order of the effect (Clausner and Gentili, 2022). All p-values are FDR-corrected (Benjamini and Hochberg, 1995).

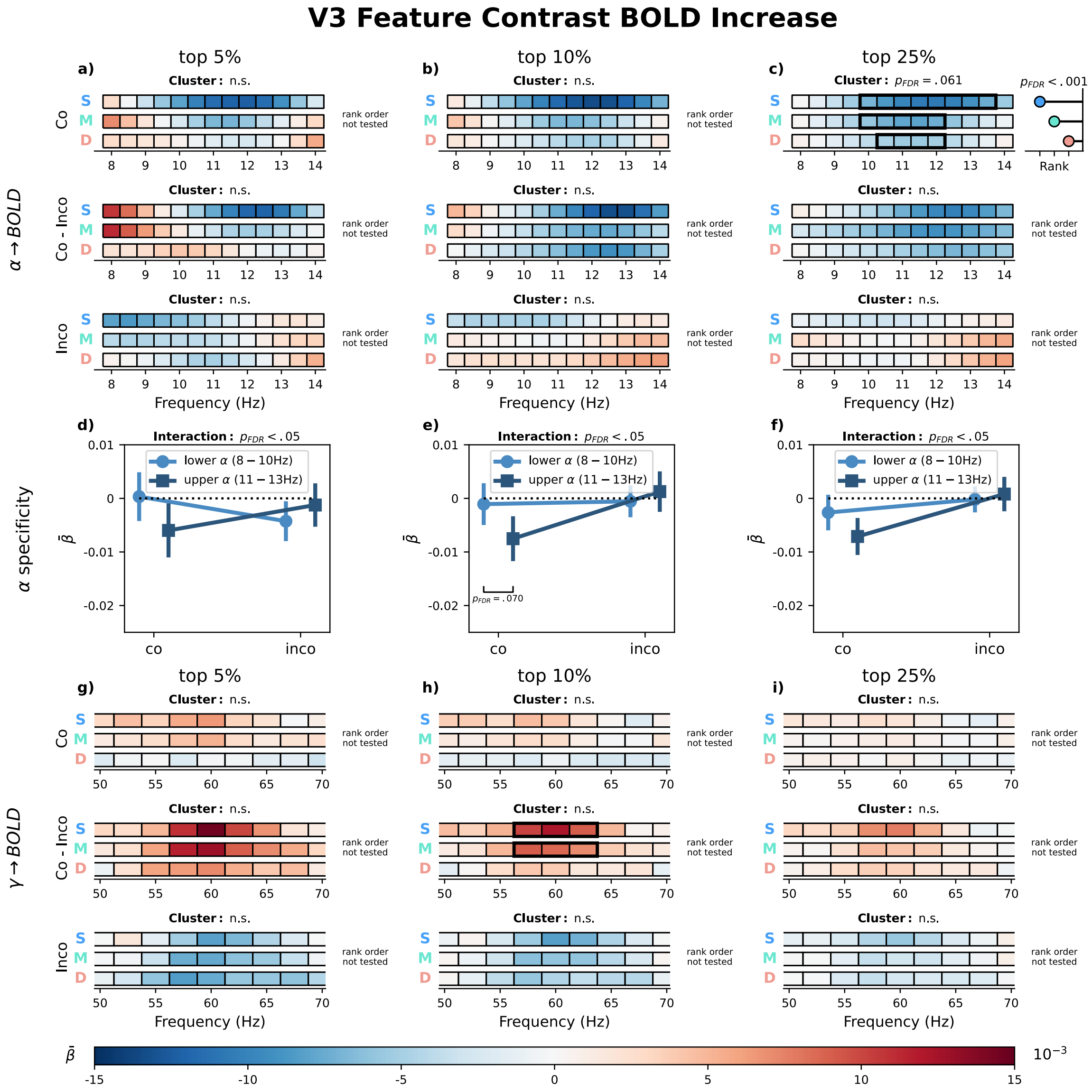


Figure 9: **V3 relationship between laminar BOLD signal and EEG for the feature-specific contrast**. Average $\bar{\beta}$ coefficients from a GLM predicting laminar BOLD signals in V3 from trial-by-trial EEG spectral power. Voxels where selected based on the first level fMRI contrast of $left - right$ oriented stimuli and their (positive) response to either stimulus orientation (feature contrast BOLD increase). Columns correspond to voxel-selection thresholds (top 5%, 10%, 25% most extreme t-values). The GLM was computed using voxels with a stronger response to one orientation over the other. EEG regressors were computed from trials of the Congruent orientation (matching the preferred voxel orientation), the Incongruent orientation (opposite). Furthermore, the difference between congruent and incongruent models has been computed (Congruent - Incongruent). $\bar{\beta}$ coefficients were weighted by laminar contribution weights, yielding estimates for superficial (S), middle (M), and deep (D) layers. **a–c**: $\alpha$-band $(8-14\text{ Hz})$ layer × frequency plots. **d–f**: average $\bar{\beta}$ (± SEM) for lower $(8-10\text{ Hz})$ and upper $(11-13\text{ Hz})$ $\alpha$ sub-bands per voxel selection, tested with a linear mixed-effects model. **g–i**: $\gamma$-band $(50-70\text{ Hz})$ layer × frequency plots. Black rectangles indicate significant clusters (Maris and Oostenveld 2007); where significant, lollipops to the right show the layer rank order of the effect (Clausner and Gentili, 2022). All p-values are FDR-corrected (Benjamini and Hochberg, 1995).

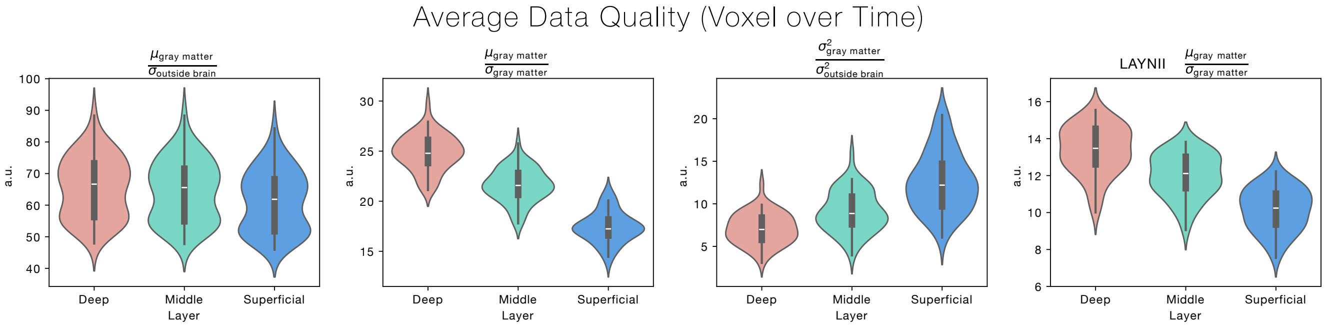


Figure 10: **Signal-to-Noise Ratio.** Several measures have been computed to estimate the temporal signal-to-noise ratio across layers, as indicated by the title of the respective sub-plot. All sub-plots except the rightmost sub-plot have been computed based on the used layering algorithm. We have included the tSNR estimate for the laminar segmentation as provided by LayNii (Huber et al., 2021), to allow for a comparison between our fraction-based approach and LayNii's approach, which assigns entire voxels to specified layers. In general, the laminar profile of tSNR estimates was unexpected, because the vascular draining effect (Markuerkiaga et al., 2016) typically leads to higher signal changes in superficial layers. However, since both functionally relevant signal components, as well as physiological noise drains towards the cortical surface, a higher tSNR for deep layers is not entirely surprising. Importantly however, the tSNR profile is not reflected in our main results, which strengthens the overall validity.

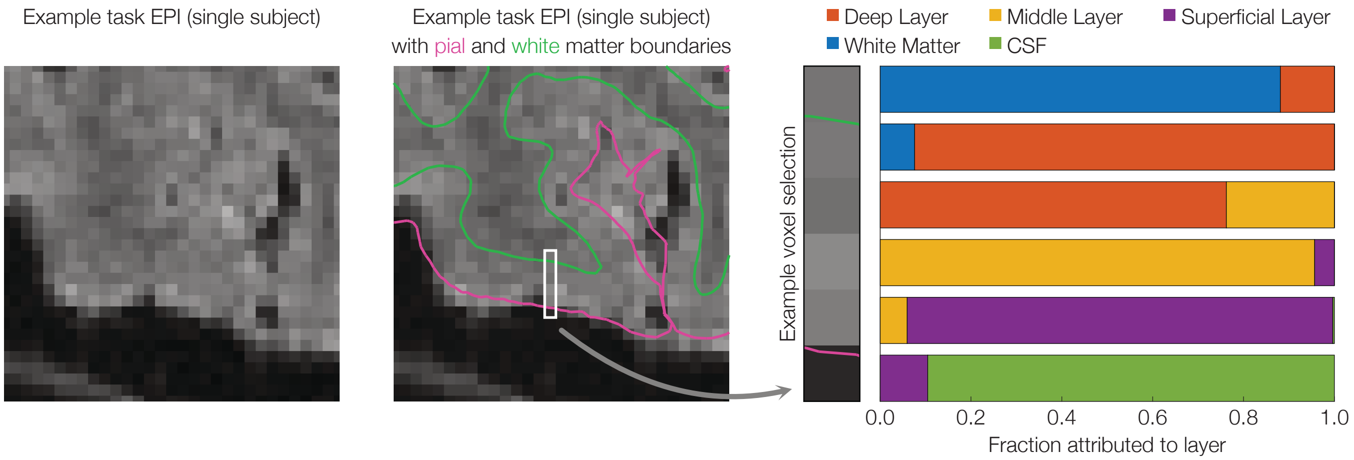


Figure 11: **EPI example with layer segmentation.** Left: a segment of a slice of a single participant's raw fMRI volume during the task. Centre: The same segment, overlaid with pial and white matter boundaries that were used for reconstructing cortical depth. The white box indicates an example voxel selection for which the respective layer attributions are shown on the Right: Each voxel was assigned a fraction by how much this voxel belongs to a specific layer.

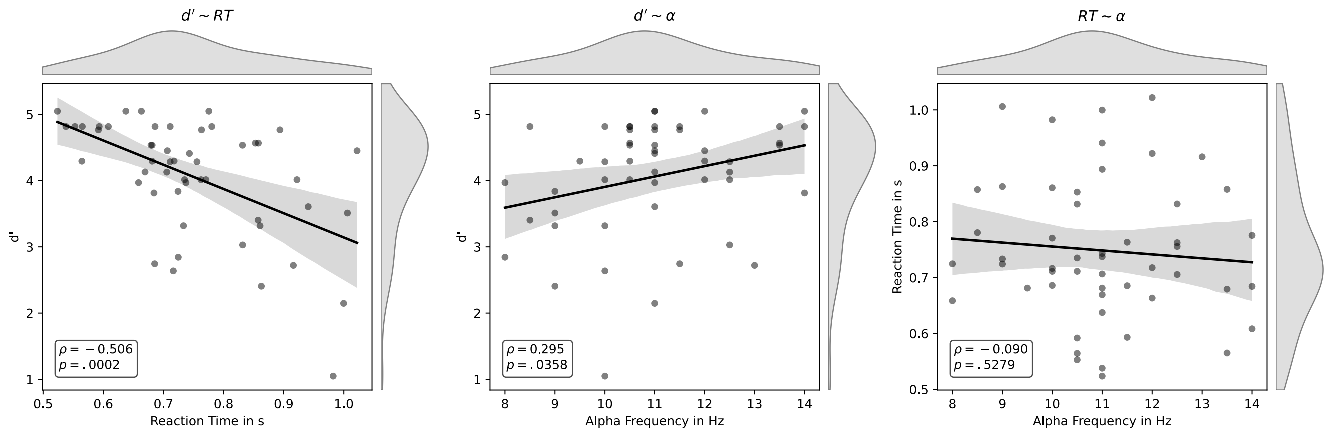


Figure 12: **Correlation of $\alpha$ Frequency and Task Performance.** Task performance for each participant has been defined as the average reaction time (RT) to correctly identified oddball stimuli, as well as d' as a measure for average response accuracy. Individual $\alpha$ frequencies have been determined for each participant based on the average over all correctly identified non-oddball trials. Within the time of interest ($0.1$ to $0.8$ s after stimulus onset) and frequencies of interest ($8$ to $14$ Hz) the frequency with the largest decrease was chosen. Afterwards, per-participant frequencies have been correlated with per-participant average RTs and d' values. We found that participants with the strongest decrease of $\alpha$ power in higher frequencies, also respond more accurately on average. No such correlation was observed between $\alpha$ frequency and RT. However, RT and d' have been found to be negatively correlated as well.

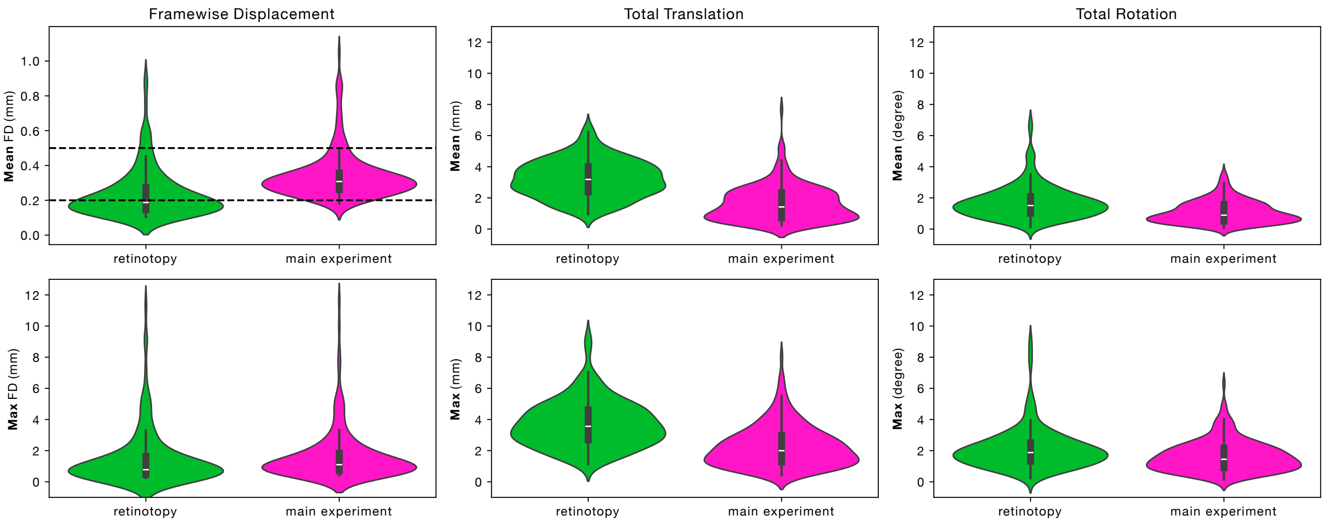


Figure 13: **Average Motion of Participants.** Average across participants for average and maximum frame-wise displacement (Power et al., 2012) using a radius of $50\ \mathrm{mm}$, total translation and total rotation within one experimental block.
